# A method to determine the double bond position in unsaturated fatty acids by solvent plasmatization using liquid chromatography-mass spectrometry

**DOI:** 10.1101/2019.12.24.875385

**Authors:** Shigeo Takashima, Kayoko Toyoshi, Takuhei Yamamoto, Nobuyuki Shimozawa

## Abstract

Fatty acids (FAs) are the central components of life: they constitute biological membranes in the form of lipid, act as signaling molecules, and are used as energy sources. FAs are classified according to their chain lengths and the number and position of carbon-carbon double bond, and their physiological character is largely defined by these structural properties. Determination of the precise structural properties is crucial for characterizing FAs, but pinpointing the exact position of carbon-carbon double bond in FA molecules is challenging. Herein, a new analytical method is reported for determining the double bond position of mono- and poly-unsaturated FAs using liquid chromatography-mass spectrometry (LC-MS) coupled with solvent plasmatization. With the aid of plasma on ESI capirally, epoxidation or peroxidation of carbon-carbon double bond in FAs is facilitated. Subsequently, molecular fragmentation occurs at or beside the epoxidized or peroxidized double bond via collision-induced dissociation (CID), and the position of the double bond is elucidated. In this method, FAs are separated by LC, modified by plasma, fragmented via CID, and detected using a time-of-flight mass spectrometer in a seamless manner such that the FA composition in a mixture can be determined. Our method enables thorough characterization of FA species with distinguishing multiple isomers, and therefore can uncover the true diversity of FAs for their application in food, health, and medical sciences.

## 1. INTRODUCTION

Fatty acids (FAs) are the central components of living organisms as they constitute biological membranes in the form of lipid, can be converted into physiologically active signal molecules (i.e. eicosanoids and lysophospholipids), and serve as energy sources [1–5]. Their chemical characteristics are primarily determined by their carbon chain length as well as the number and position of carbon-carbon double bonds. Saturated fatty acids (SFAs) do not contain carbon-carbon double bonds and their chemical stability is suitable for cellular energy storage. In contrast, unsaturated FAs contain one or more carbon-carbon double bonds and are relatively unstable compared to SFAs, especially when multiple double bonds are present. Unsaturated FAs with a single carbon-carbon double bond are referred to as mono-unsaturated fatty acids (MUFAs), while those with multiple carbon-carbon double bonds are collectively called poly-unsaturated fatty acids (PUFAs). Unsaturated FAs can be further categorized into several groups, such as *ω*-3, *ω*-6, and *ω*-9 family FAs, where each family is distinguished by the position of the first double bond in relation to the omega carbon. The number and position of double bonds in FAs dictate their biological activities and are considered to be important subjects for human health [2,6,7].

Gas chromatography-mass spectrometry (GC-MS) is traditionally used to analyze FAs. To determine the position of carbon-carbon double bonds, derivatization of unsaturated FAs at the double bonds, such as with dimethyl disulfide as a derivatization agent, is often applied. The derivatized unsaturated FAs are ionized using electron ionization (EI) in GC-MS. The high energy of EI enables to cleave the modified bond to elucidate the diagnostic fragment ions from which the position of the double bond can be determined [8–11]. Liquid chromatography-mass spectrometry (LC-MS) complements GC-MS in many applications including FA analysis, and it is adequate for very-long-chain FAs (FAs with 22 or more carbons) that have poor volatility in GC-MS [12]. The determination of the position of carbon-carbon double bond in unsaturated FAs using LC-MS, however, remains challenging. Electrospray ionization (ESI) and atmospheric pressure chemical ionization (APCI), both often used in LC-MS, are the methods involving soft ionization; so, fragmentation does not occur effectively by itself. Subsequently, collision-induced dissociation (CID) is used to fragment the ionized FAs. However, it usually does not provide the informative fragment ions to determine the double bond position. To deduce the informative fragment ions by CID, modification of FAs on carbon-carbon double bonds or the terminal carboxyl group is required. For example, acetonitrile-related adducts of FA methyl esters produce diagnostic fragment ions from which the double bond position can be determined [13,14]. Epoxidation of the double bond of FAs by low-temperature plasma successfully produces the informative diagnostic fragment ions from free FAs and lipid-conjugated FAs after CID fragmentation [15,16]. Paper-spray ionization followed by CID also successfully epoxidize FAs and produce the same type of diagnostic fragment ions to identify the position of carbon-carbon double bonds [17]. Paternó–Büchi reaction with acetone also modifies the double bond, which subsequently produces diagnostic fragment ions via CID [18–21]. Derivatization with *N*-(4-aminomethylphenyl)pyridinium (AMPP) at the terminal carboxy group also provides the informative fragment ions for carbon-carbon double bond [22,23]. With these methods one can deduce the exact position of the carbon-carbon double bond of unsaturated FAs.

We previously reported a method for analyzing a range of FA species in biological samples in a single assay by using reverse-phase LC-MS [12,24], where FAs were distinguished by their chain lengths and number of double bonds. Using this method, approximately 50 FA species could be distinguished in control human fibroblasts and >100 species in patient fibroblasts with Zellweger syndrome (ZS), wherein the biosynthesis of peroxisomes is impaired and FA metabolism is altered. Although we could estimate the relative position of the double bond among FA isomers, the exact position of double bonds in unsaturated FAs could not be determined.

In the present study, we established a method that can determine the exact position of the carbon-carbon double bonds in unsaturated FAs by using conventional LC-MS without specialized setups. Herein, we induced epoxidation and peroxidation of the carbon-carbon double bond of unsaturated FAs facilitated by solvent plasmatization as corona discharge. Subsequent CID produced the diagnostic fragment ions for determining the double bond position. Peroxidation, but not epoxidation, of PUFA was found to be required for producing the informative diagnostic fragment ions. We applied our method to analyze FAs from biological samples. In combination with our previous method, the current method enables full characterization of a FA species and through profiling of the FA composition in biological samples or in food resources.

## 2. Materials and methods

### 2.1. Reagents

The FA standards used in this study were as follows: *cis*-6-octadecenoic acid (C18:1 *ω*-12, petroselinic acid; Sigma-Aldrich, St. Louis, MO, USA, #P8750), *cis*-9-octadecenoic acid (C18:1 *ω*-9, oleic acid; Cayman Chemical, Ann Arbor, MI, USA, #90260), *cis*-11-octadecenoic acid (C18:1 *ω*-7, *cis*-vaccenic acid; Matreya LLC, State College, PA, USA, #1266), ^13^C-UL-oleic acid (^13^C-labeled oleic acid; Martek Isotopes, Olney, MD, USA), all *cis*-9, 12, 15-octadecatrienoic acid (C18:3 *ω*-3, α-linolenic acid; Cayman Chemical, #90210), all *cis*-6, 9, 12-octadecatrienoic acid (C18:3 *ω*-6, γ-linolenic acid; Cayman Chemical, #90220), all *cis*-5, 11, 14-eicosatrienoic acid (20:3 *ω*-6, sciadonic acid; Cayman Chemical, # 10009999), all *cis*-8, 11, 14-eicosatrienoic acid (20:3 *ω*-6, dihomo-γ-linolenic acid; Cayman Chemical, #90230), all *cis*-5, 8, 11, 14, 17-eicosapentaenoic acid (20:5 *ω*-3, EPA; Cayman Chemical, #90110.1), all *cis*-5, 8, 11, 14-eicosatetraenoic acid (20:4 *ω*-6, arachidonic acid, ARA; Cayman Chemical, #90010), all *cis*-4, 10, 13, 16-docosatetraenoic acid (C22:4 *ω*-6, *cis*-4 DTA, Cayman Chemical, #10007289), all *cis*-4, 7, 10, 13, 16, 19-docosahexaenoic acid (22:6 *ω*-3, DHA, Tokyo Chemical Industry/TCI, Tokyo, Japan, #D2226). The FA standards were dissolved in a solution containing two volumes of chloroform and one volume of methanol with 0.05% (w/v) of 2,6-di-*t*-butyl-*p*-cresol (butylated hydroxytoluene; Nacalai Tesque, Kyoto, Japan, #11421-92) and stored at −20 °C until use. Other reagents used were: Deuterium oxide (D_2_O, FUJIFILM Wako Pure Chemical Corp., Osaka, Japan, #049-34242), Water-^18^O (H_2_^18^O, Taiyo Nippon Sanso Corp., Tokyo, Japan, #F03-0027), high-performance liquid chromatography (HPLC)-grade *tert*-butyl methyl ether (*t*-BME, Sigma-Aldrich, #34875), HPLC-grade acetonitrile (FUJIFILM Wako, #019-08631), HPLC-grade acetone (Wako, #014-08681), HPLC-grade 1 mM ammonium acetate (FUJIFILM Wako, #018-21041), and 10% ammonia solution (FUJIFILM Wako, #013-17505).

### 2.2. Liquid chromatography-mass spectrometry

A Waters Acquity ultra-performance liquid chromatography system equipped with an auto sampler and reverse-phase column with thermal control was used. A Waters BEH C_8_ column (2.1 × 50 mm, particle size 1.7 μm, pore size 130 Å, #186002877) preceded by a Waters BEH C_8_ VanGuard Pre-Column (2.1 × 5 mm, particle size 1.7 μm, pore size 130 Å, #186003978) was used. Aqueous mobile phase A consisted of purified water containing 1 mM (0.0077% w/v) ammonium acetate and 5.78 mM (0.01% w/v) ammonia. The organic mobile phase B was 100% acetonitrile. The flow rate was 0.1 mL/min and the following linear gradient was applied (indicated content of mobile phase B): 0 to 50 min, 20% to 95% linear increase; 50 to 52.5 min, hold at 95%; 52.5 to 55 min, hold at 20%. Mass spectrometry was performed using a Waters XevoQTof MS system, which is a time-of-flight mass spectrometer preceded by a quadrupole and collision chamber. The mobile phase was plasmatized by charging high voltage on the ESI capillary to cause corona discharge. The samples were applied to MS from the LC system or via direct infusion channel without LC separation. The MS settings were as follows: capillary voltage on negative ESI, 1.0 kv for normal MS assay and 3.6 to 4.2 kV for plasmatization; sampling cone, 56 (arbitrary value); extraction cone, 4.0 (arbitrary value); source temperature, 125 °C; desolvation temperature, 350 °C; cone gas flow, 60 L/h; and desolvation gas flow, 1000 L/h. Argon gas was used for collision-induced dissociation (CID). We removed lockspray baffle because it interferes stable plasma generation at the ESI tip so that lock mass was not used throughout the analysis. The obtained data were analyzed by Waters Masslynx software and ACD ChemSketch was used to draw the chemical structures.

### 2.3. Direct infusion assay

For direct infusion assay, the FA standards were dissolved in 100% acetonitrile and directly introduced in MS via an infusion port and mixed with LC solvents A and B before infusion. The same MS settings as above were used. In some experiments, the FA standards were dissolved in a mixed solution of H_2_^18^O and acetonitrile or D_2_O and acetonitrile, and were analyzed without mixing with LC solvent.

### 2.4. Purification of fatty acids from human fibroblasts

Total FAs were purified from control human skin fibroblasts, NB1RGB, obtained from the National Bio-Resource Project (NBRP) of the Ministry of Education, Culture, Sports, Science, and Technology (MEXT), and from fibroblasts of a patient with ZS with *PEX2* mutation (F-12 line) [12]. A pellet of approximately 1 × 10^6^ cells was dissolved in 400 μL of acetonitrile and 50 μL of 5 M hydrochloric acid in a glass tube. The sample was lysed by vortexing for 1 min and incubated at 100 °C for 1 h in an oil bath. After cooling to room temperature (approximately 20°C), 800 μL of *t*-BME, 100 μL of methanol containing ^13^C-labeled oleic acid (internal standard; 1 μg/mL) and 400 μL of purified water were added, and vortexed for 1 min. Phase separation was achieved by a centrifugation at 200 ×g for 5 min, and the upper organic phase was collected. Subsequently, 800μL of water was added and the sample was vortexed again for 1 min. After phase separation at 200×g for 5 min, the upper organic phase containing FAs was collected and placed in a glass sample vial with a polytetrafluoroethylene (PTFE)-lined cap (GL Sciences, Tokyo, Japan, #1030-51023 and #1030-45260). The organic phase was evaporated under a stream of nitrogen gas and the recovered FAs were reconstituted in 100 μL of acetone and subjected to LC-MS analysis.

This study was approved by the Ethical Committee of the Graduate School of Medicine, Gifu University (permission number: 29-286).

## 3. Results and discussion

### 3.1. Solvent plasmatization facilitates epoxidation and peroxidation of unsaturated fatty acids for double bond position determination

Zhao et al reported that epoxidation of unsaturated FAs by low-temperature plasma and subsequent fragmentation using ESI-MS can determine the position of carbon-carbon double bond [15]. We tried a similar approach using LC-ESI-MS and found that unsaturated FAs can be epoxidized and peroxidized by plasmatization of the LC solvent. When excessively high electric voltage was applied at the tip of the ESI capillary, plasmatization of the solvent occurred as corona discharge (Figure 1A). Epoxidation of oleic acid (C18:1 *ω*-9, *cis*-9) was first attempted with direct infusion assay (100-200 pmol/min with 0.1 mL/min of solvent; isocratic 50% B). At a normal capillary voltage on negative ESI (1.0 kv), ionized oleic acid ([M-H]–, *m/z* 281.25) was detected (Supplementary Figure S1A). When a high voltage (≥3.4 kV) was applied to the ESI capillary, plasmatization of the solvent was induced at the capillary tip, and simultaneously, the epoxidized oleic acid ([M-H+O]–, *m/z* 297.24) was detected in the mass spectrum (Figure 2A, left panel; Figure S1B). The molecule was fragmented by CID and produced fragment ions at *m/z* 171.10 and 155.11 (Figure 2A, right panel; Figure 2D), consisting of the fragments containing the alpha carbon (hereafter “alpha fragments”), from which the double bond position can be determined [15,17]. The same protocol was attempted using *cis*-vaccenic acid (C18:1 *ω*-7, *cis*-11) and petroselinic acid (C18:1 *ω*-12, *cis*-6), which have the same chain length and number of double bonds, but different double bond positions compared to those in oleic acid. Following epoxidation and CID, similar results were obtained while producing the diagnostic fragments for the determination of the double bond position (Figure 2B-D). Thus, solvent plasmatization as corona discharge was found to be useful for the epoxidation of FAs and this method is referred to as plasma-ESI-MS. We next combined this method with LC analysis. The entire scheme of the analysis, LC-plasma-ESI-MS, is depicted in Figure 1B; and the diagnostic fragment ions for determining the position of carbon-carbon double bond in MUFAs and PUFAs found in the above and following experiments are shown in Figure 1C and 1D.

**Figure 1.**
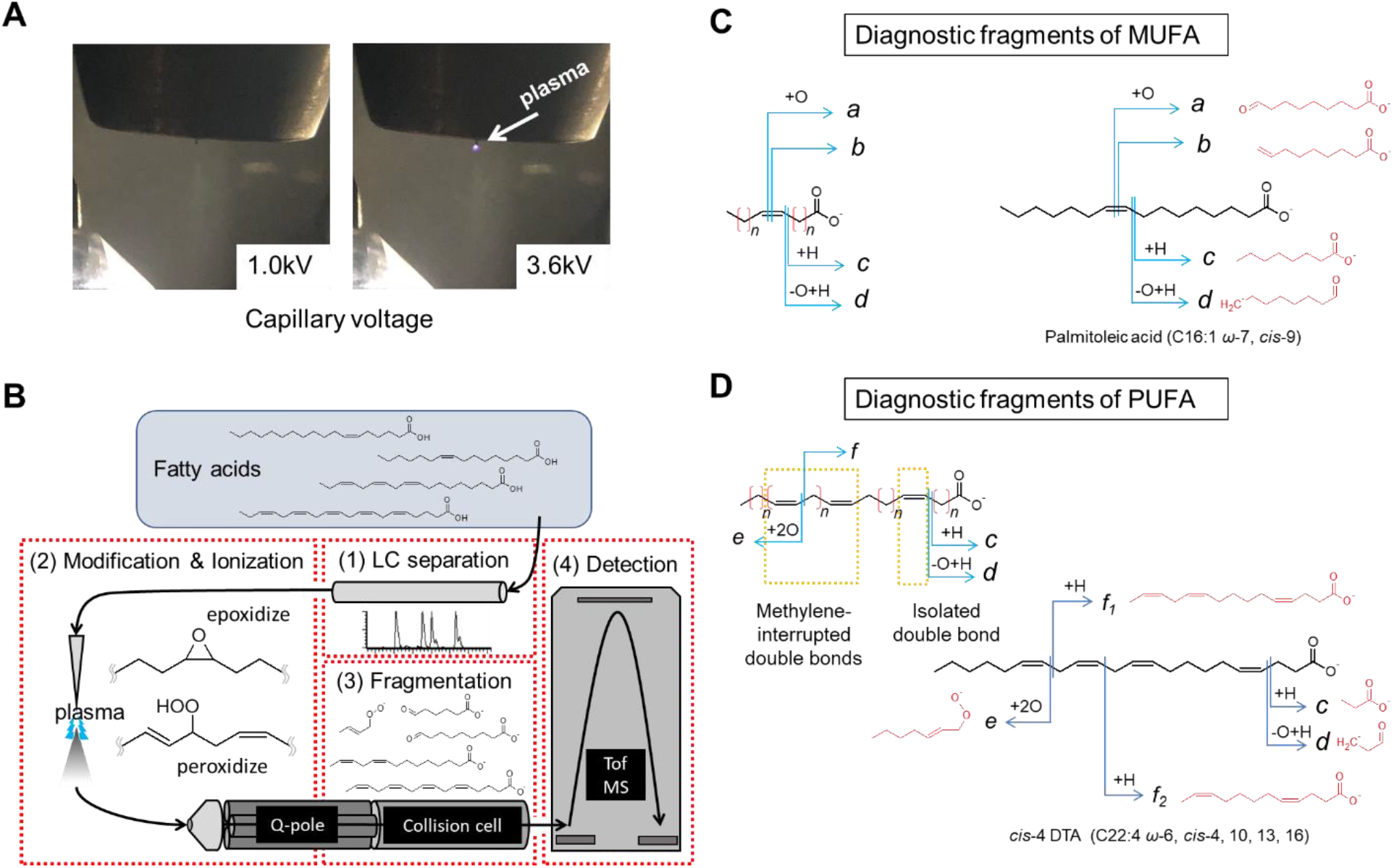
Summary of the method to determine the position of carbon-carbon double bonds in unsaturated FAs using LC-MS. (A) Plasmatization of the LC solvent on ESI probe at different electrical voltages. At a normal voltage (left, 1.0kV), no plasmatization occurs, whereas plasmatization occurs at high voltage (right, 3.6 kV) at the tip of the ESI capillary (arrow). (B) A schematic of the LC-plasma-ESI-MS assay developed in this study. Q-pole: quadrupole, Tof MS: time-of-flight mass spectrometer. (C, D) Fragmentation patterns of a representative MUFA (C) and PUFA (D). Generalized fragmentation patterns are shown on the left and real examples are shown on the right. Diagnostic fragments (fragment *a* to fragment *f*) are indicated with expected molecular structures.

**Figure 2.**
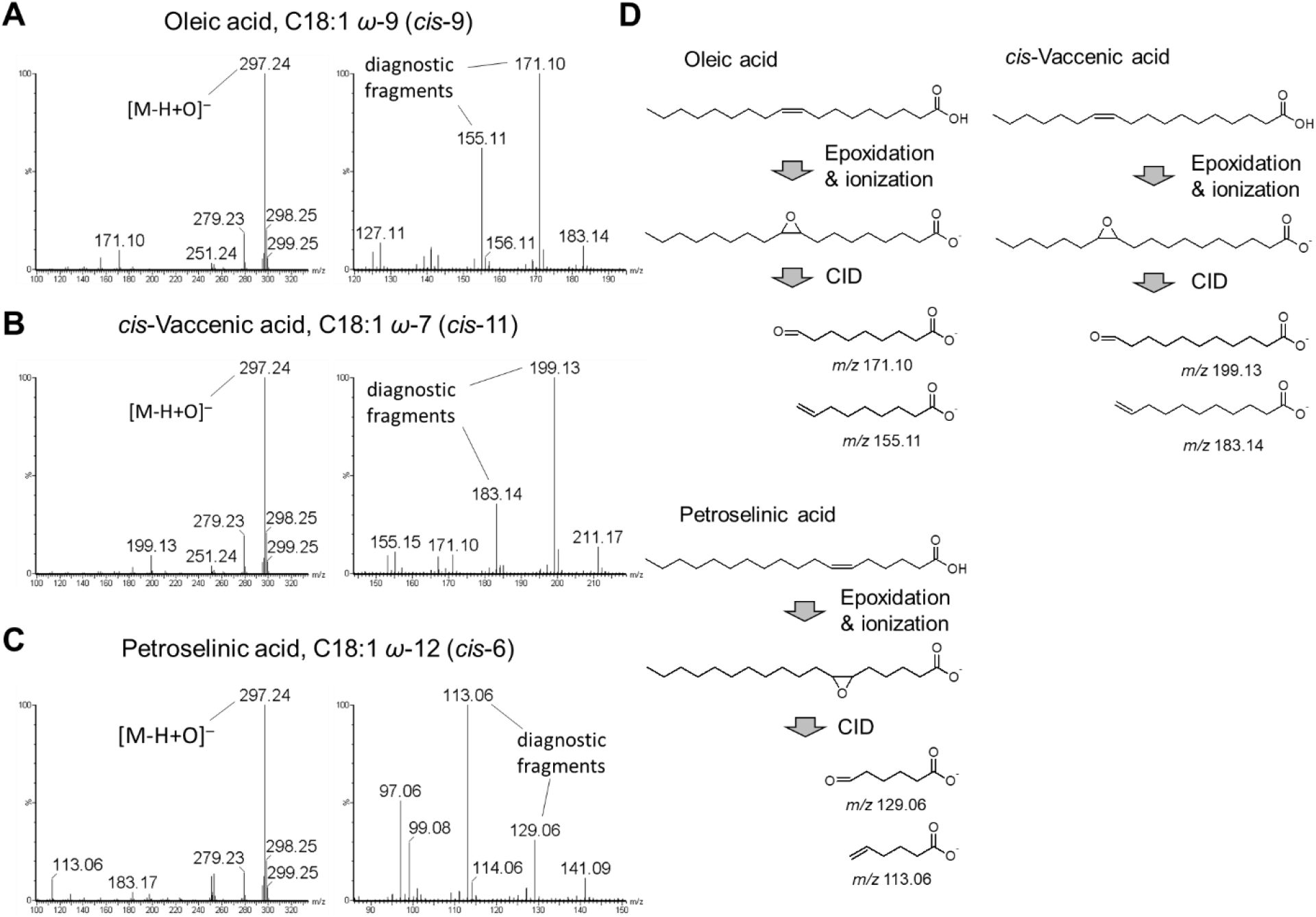
Structural determination of octadecenoic acids (C18:1). (A-C) Analysis of oleic acid (A), *cis*-vaccenic acid (B), and petroselinic acid (C) using a direct infusion assay. The parent ion ([M-H+O]–, *m/z* 297.24; marked in the left panels) and diagnostic fragment ions (right panels) of three C18:1 species are shown. (D) Flow of the diagnostic fragment formation. Throughout the manuscript, *y*-axis of the graphs indicates relative intensity and *x*-axis indicates either *m/z* (in spectral charts) or retention time (in mass chromatograms).

### 3.2. Determination of the double bond position of MUFAs by LC-MS

We analyzed octadecenoic acids (C18:1 FAs) with LC-plasma-ESI-MS. In our previous study [24], the proximity of the double bond to the omega carbon was found to be the major determinant of retention time of unsaturated FAs in reverse-phase LC. *Cis*-vaccenic acid (*ω*-7) elutes from the LC column first, oleic acid (*ω*-9) elutes second, and petroselinic acid (*ω*-12) elutes last [24]. Here, this was also confirmed by determining the exact position of the double bond. C18:1 FAs were injected independently and simultaneously as a mixed sample. In the independent injection, each C18:1 species yielded a sharp peak at a distinct retention time (Figure S2A). The epoxidized form of each C18:1 species was detected, and by CID, the diagnostic fragment ions were produced (Figure S2B and S2C; fragment *a* and fragment *b* in Figure 1C). In the mixture injection, peaks were imperfectly separated (Figure 3A-C). The epoxidized form was detected for all peaks. Upon CID, the fragment ions of the epoxidized *cis*-vaccenic acid were observed at the earliest retention time at the first half of the first peak (Figure 3A), the fragments of epoxidized oleic acid were observed at the second half of the first peak (Figure 3B), and the fragments of epoxidized petroselinic acid were observed in the second peak (Figure 3C). Epoxidized *cis*-vaccenic acid showed fragments at *m/z* 199.13 and 183.14 with a peak position at 17.82 min (Figure 3A). The fragments for the epoxidized oleic acid, *m/z* 171.10 and 155.11, showed peaks at 18.24 and 18.14 min, respectively (Figure 3B). The fragments for the epoxidized petroselinic acid, *m/z* 129.06 and 113.06, showed a peak at 18.66 min (Figure 3C). These time differences confirmed the elution order of C18:1 FA species: *cis*-vaccenic acid, oleic acid, and then petroselinic acid. The above data confirmed our previous observations that showed the order of elution timing of unsaturated FAs correlated with the proximity of the double bond to the omega carbon.

**Figure 3.**
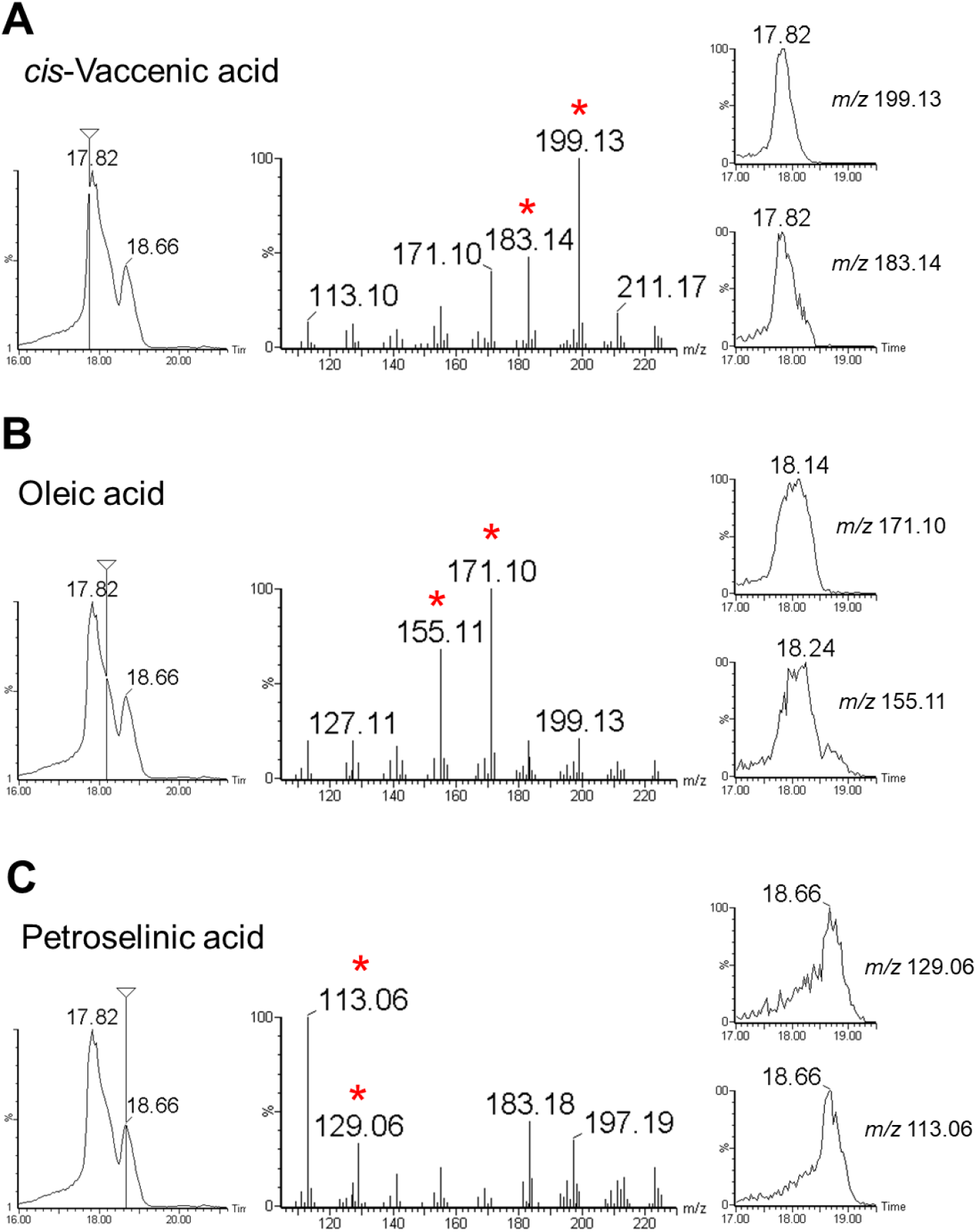
Determination of the position of double bonds in C18:1 FAs using LC-plasma-ESI-MS. (A–C) Oleic acid, *cis*-vaccenic acid, and petroselinic acid were simultaneously injected and analyzed via LC-MS with plasma-mediated epoxidation. Total ion chromatogram from the MS/MS assay, with peak top time is shown in the left panels. The time points where the fragment spectra were extracted are indicated with vertical bars. The extracted spectra are shown in the middle panels and diagnostic fragment ions are indicated by asterisks. The tandem mass chromatograms of the diagnostic fragment ions are shown in the right panels with the peak top time indicated.

We found that FAs can also be peroxidized by the plasmatization of the solvent, and the fragmentation spectra of the peroxidized FAs are also informative for determining the position of the double bonds. On solvent plasmatiozation, peroxidized oleic acid (*m/z* 313.24) was found simultaneously with the epoxidized oleic acid, though the intensity is lower than the epoxidized form (Figure S1B). Upon CID, peroxidized oleic acid produced the same fragment ions as the epoxidized form (Figure S3A and S3B), such as the alpha fragments at *m/z* 171.10 and 155.11. Additional fragment ions were also observed at *m/z* 143.11 and 127.11 corresponding to the alpha fragments with and without terminal oxygen (fragment *c* and fragment *d* in Figure 1C). Minor fragment ions such as peroxidized alpha fragment at *m/z* 201.11, and the fragment at *m/z* 141.13 corresponding to the omega carbon-side fragment (omega fragment) were also detected (Figure S3A and S3B). Similar results were obtained for the analysis of peroxidized *cis*-vaccenic acid and petroselinic acid (Figure S3C-F). These additional fragments yielded positional information of the double bond, and similar fragments were informative for determining the position of double bonds in PUFAs, which will be discussed in subsequent sections.

### 3.3 Determination of the position of double bonds in PUFAs

We next examined if the plasma-mediated modification of PUFAs could yield any informative fragment spectra for determining the position of carbon-carbon double bonds. Two types of octadecatrienoic acids (C18:3), α-linolenic acid (C18:3 *ω*-3, *cis*-9, 12, 15) and γ-linolenic acid (C18:3 *ω*-6, *cis*-6, 9, 12) were first examined by LC-plasma-ESI-MS. They were successfully epoxidized or peroxidized by solvent plasmatization, and upon CID, the epoxidized parent ions (*m/z* 293.22) yielded highly prominent fragment spectra at *m/z* 71.05 in α-linolenic acid and *m/z* 113.10 in γ-linolenic acid (Figure 4A and 4B, left panels). These spectra corresponded to the omega fragments epoxidized at the proximal double bond to the omega carbon. These type of fragmentation spectra always appeared in extremely high intensities in PUFAs and can be used for determining the double bond that is closest to the omega carbon for their categorization into *ω*-n families. When peroxidized α-linolenic acid and γ-linolenic acid (*m/z* 309.21) were fragmented, ions at *m/z* 87.05 and 129.09, respectively, were detected in high intensities, and were peroxidized omega fragments at the proximal double bond to the omega carbon (Figure 4A and B, right panels). The alpha fragments at *m/z* 223.17 and 181.12, which are the counterpart fragments of the above omega fragments, were also detected in high intensities (Figure 4A and B, right panels). From the omega and alpha fragments, the positions of the first and second double bonds can be determined. Similar alpha fragments were observed at the second and third double bonds (*m/z* 183.14 and 141.09 for α- and γ-linolenic acid, respectively; Figure 4A and B). From these diagnostic fragments, the positions of all the three double bonds can be determined. The fragmentation yielding this type of alpha fragments only effectively occurs when the neighboring double bonds are separated by a single methylene group (hereafter referred to as “methylene-interrupted double bonds” Figure 1D) and not by multiple methylene groups (hereafter referred to as “isolated double bond”; Figure 1D). In addition, there were multiple minor fragment ions that could support the positional information of double bonds as shown in Figures S4 and S5). Because the fragmentation of the peroxidized form of PUFA always yields highly intense spectra of the diagnostic fragment ions, peroxidized forms are primarily used to determine the positions of double bonds in PUFAs and their corresponding epoxidized forms are used to support the obtained results.

**Figure 4.**
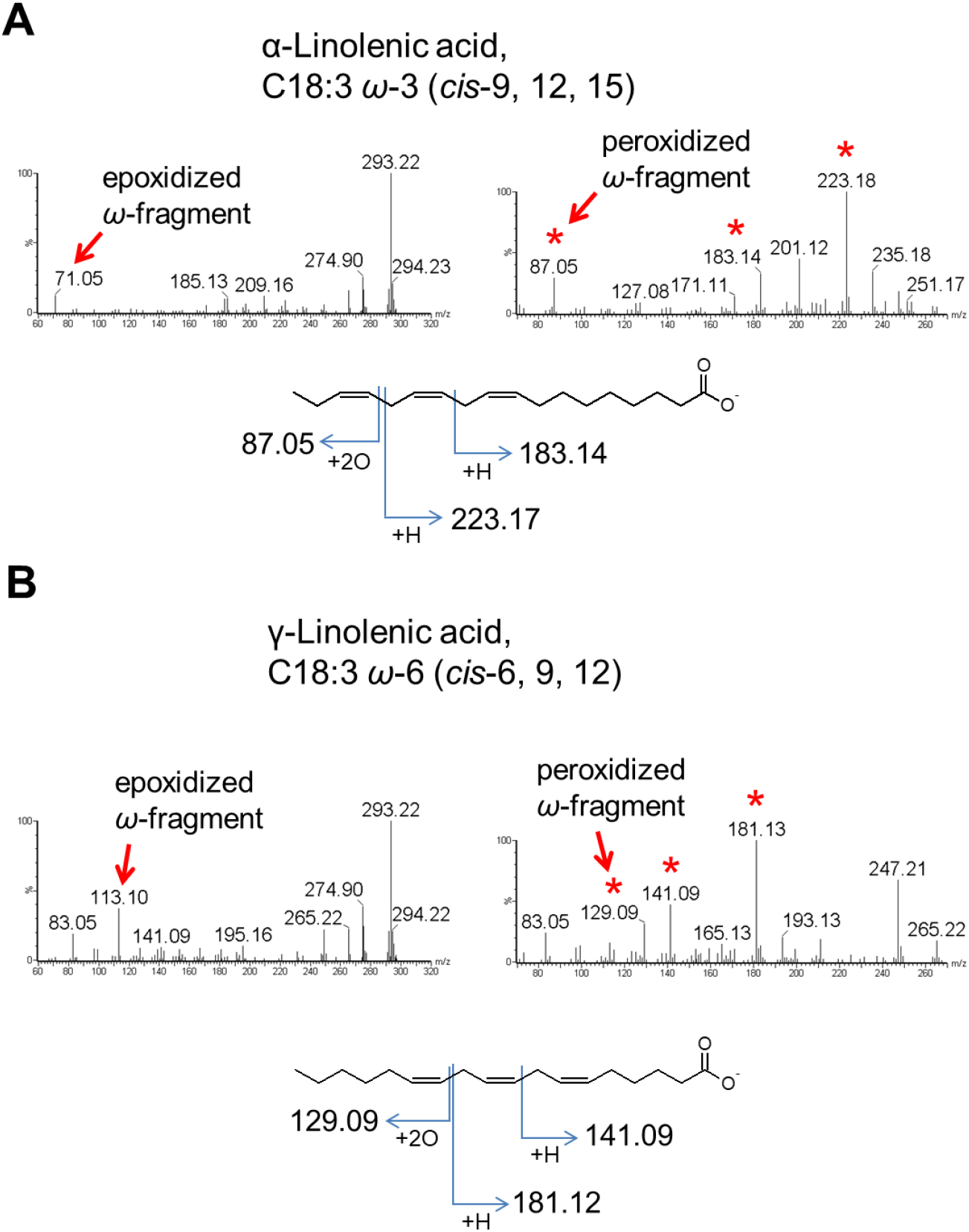
Structural determination of C18:3 FAs. (A, B) α- and γ-linolenic acid were fragmented after epoxidation (left panels) or peroxidation (right panels). The epoxidized and peroxidized fragments at the proximal double bond to the omega carbon are indicated by arrows and the diagnostic fragment ions from peroxidized C18:3 FAs are indicated by asterisks. The graphical fragmentation patterns for diagnostic fragment ions after peroxidation are also shown.

Two types of eicosatrienoic acid (C20:3), sciadonic acid (C20:3 *ω*-6 *cis*-5, 11, 14) and dihomo-γ-linolenic acid (C20:3 *ω*-6 *cis*-8, 11, 14), were examined. Dihomo-γ-linolenic acid contains three methylene-interrupted double bonds, whereas sciadonic acid contains two methylene-interrupted double bonds and one isolated double bond (Figure 5C). From the fragmentation pattern of the peroxidized form, dihomo-γ-linolenic acid showed a similar fragmentation pattern as that of the C18:3 fatty acids described above (Figure 5A and C). In contrast, sciadonic acid yielded the same fragmentation pattern only from its methylene-interrupted double bonds (*cis*-11 and *cis*-14 double bonds) (Figure 5B and C). The isolated double bond at *cis*-5 position behaved in a similar manner as a MUFA double bond, producing common alpha fragments (Figure 5B and C, Figure S3), enabling the determination of the position of the isolated double bond. The same fragmentation pattern was observed with the isolated double bond of *cis*-4 DTA, which contains three methylene-interrupted double bonds and an isolated double bond (see below). Therefore, the double bond positions in PUFAs can be determined by the fragment ions specific to the methylene-interrupted and isolated double bonds.

**Figure 5.**
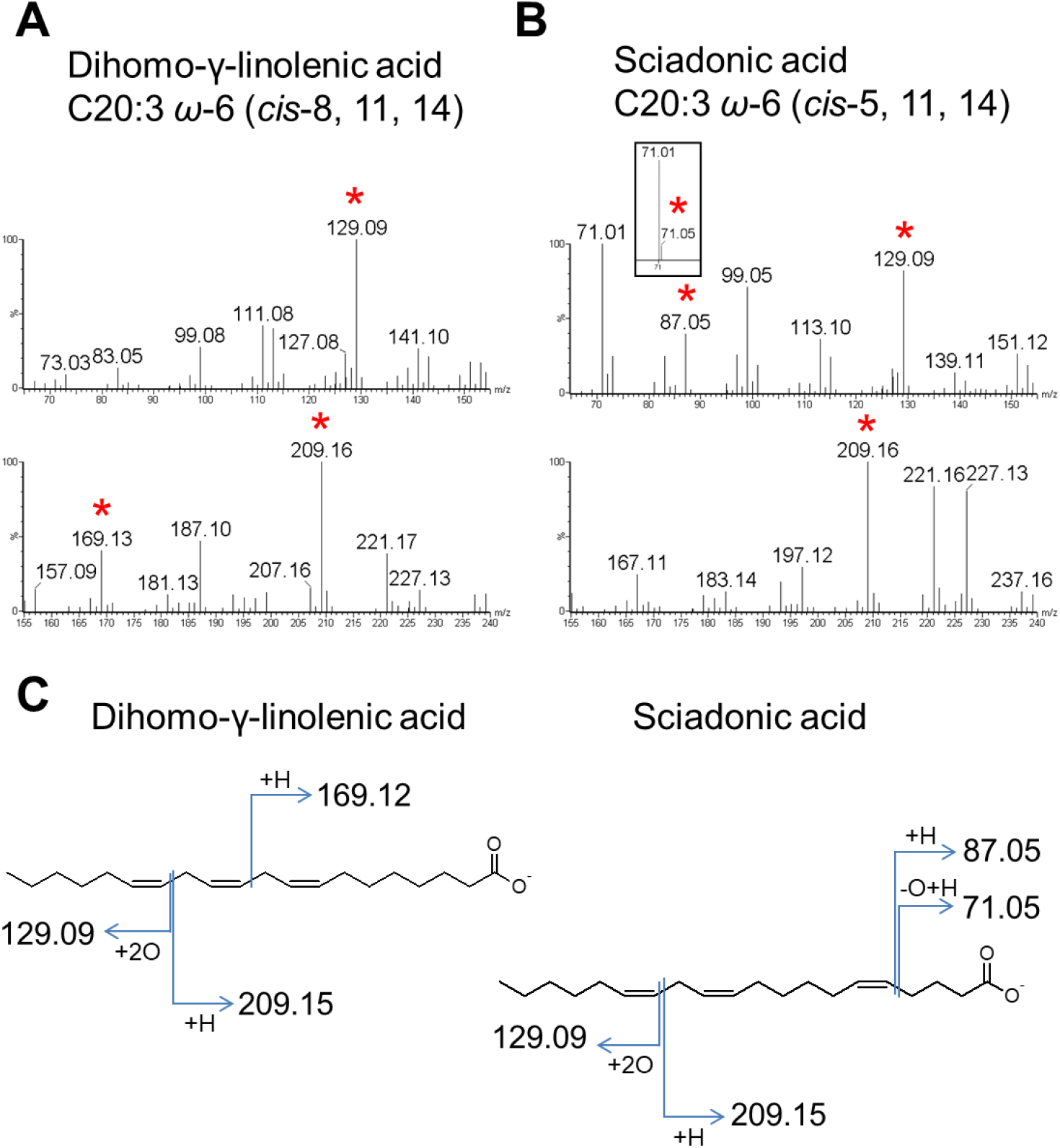
Structural determination of C20:3 FAs. (A, B) Dihomo-γ-linolenic acid and sciadonic acid were peroxidized and fragmented. The diagnostic fragment ions are indicated by asterisks. It should be noted that common fragment ions originate from the identical double bonds while specific fragment ions originate from the distinct double bonds. (C) Graphical fragmentation patterns for C20:3 FAs are shown.

In PUFAs with four or more double bonds, diagnostic alpha and omega fragments were more prominent, while the minor fragments decreased in intensity. As the PUFAs shown above, the methylene-interrupted double bonds in ARA (C20:4 *ω*-6, *cis*-5, 8, 11, 14) (Figure 6A), *cis*-4 DTA (C22:4 *ω*-6, *cis*-4, 10, 13, 16) (Figure 6B), EPA (C20:5 *ω*-3, *cis*-5, 8, 11, 14, 17) (Figure 6C), and DHA (22:6 *ω*-3, *cis*-4, 7, 10, 13, 16, 19) (Figure 6D) yielded the same type of diagnostic omega and alpha-fragments (fragments *e* and *f*s in Figure 1D). The isolated double bond in *cis*-4 DTA can be positioned using the common alpha fragments with MUFA (*m/z* 73.03 and 57.03; fragments *c* and *d* in Figure 1D). The above data demonstrate that the double bond position of a wide range of PUFA species can be determined by peroxidation and CID fragmentation with the common fragmentation criteria.

**Figure 6.**
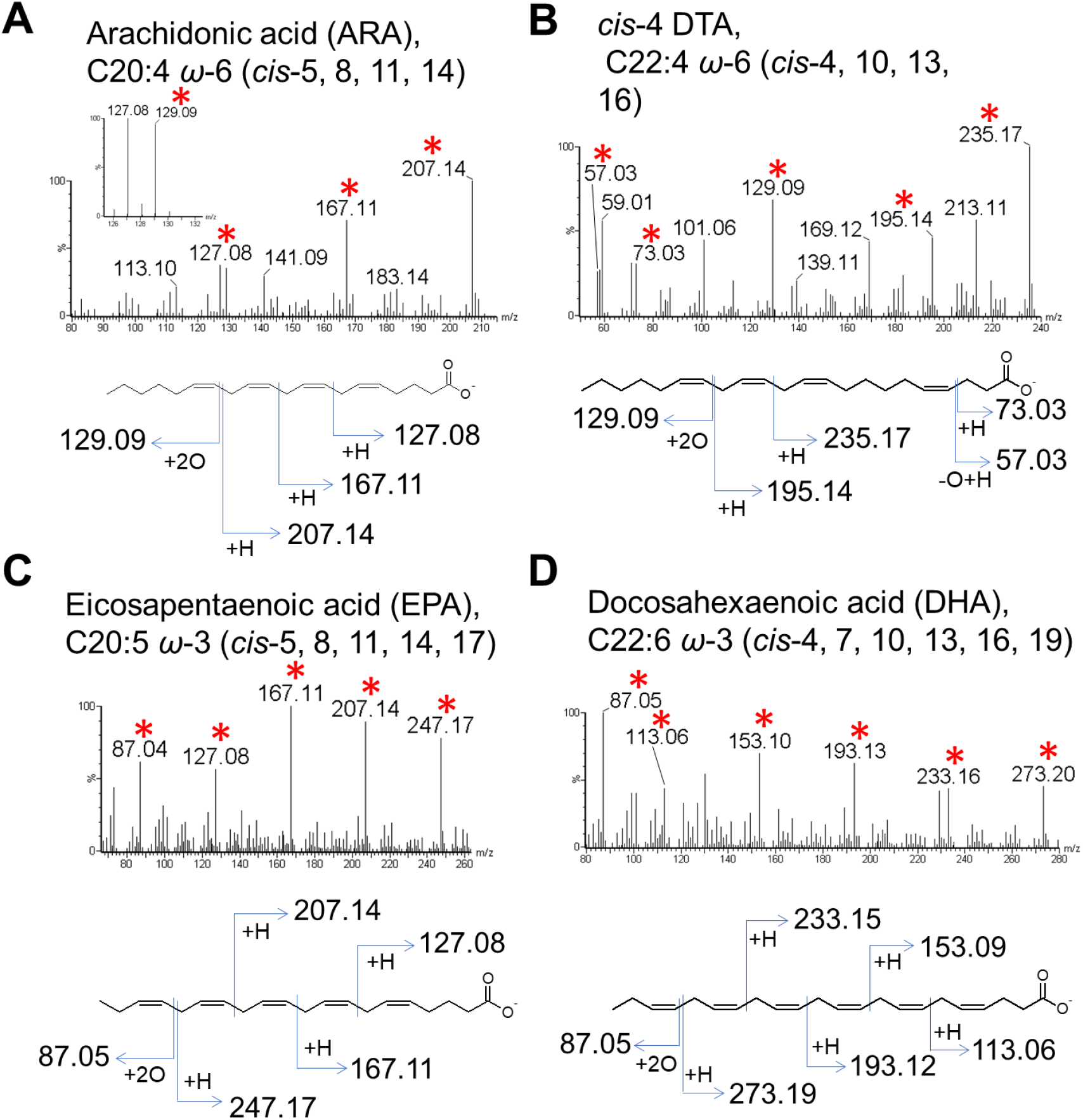
Structural determination of the PUFAs with four or more double bonds. (A, B) Peroxidized arachidonic acid (ARA) (A) and *cis*-4 DTA (B) were fragmented by CID. (C, D) Highly unsaturated FA, eicosapentaenoic acid (EPA), containing five double bonds (C), and docosahexaenoic acid (DHA) with six double bonds (D), were peroxidized and fragmented. The diagnostic ions are indicated by asterisks.

### 3.4. Analytical performance

The analytical performance of this method was evaluated. Standard solutions containing single FA species, such as palmitoleic acid (C16:1, *cis*-9), *cis*-vaccenic acid (C18:1, *cis*-11), dihomo-γ-linolenic acid (C20:3, *cis*-8, 11, 14), *cis*-4 DTA (C22:4 *cis*-4, 10, 13, 16), and EPA (C20:5 *cis*-5, 8, 11, 14, 17), were quantified. Calibration functions were calculated for all the diagnostic fragment ions, and good linearity was confirmed in both MUFA and PUFA species (Table 1, Figure S6). We used the diagnostic fragment ion giving the least intensity to calculate the limit of detection (LOD) and limit of quantification (LOQ) of the corresponding FA species. The calculated LOD was 6 μM to 15 μM, and LOQ was 19μM to 45 μM among examined FA species (Table 1).

**Table 1.**
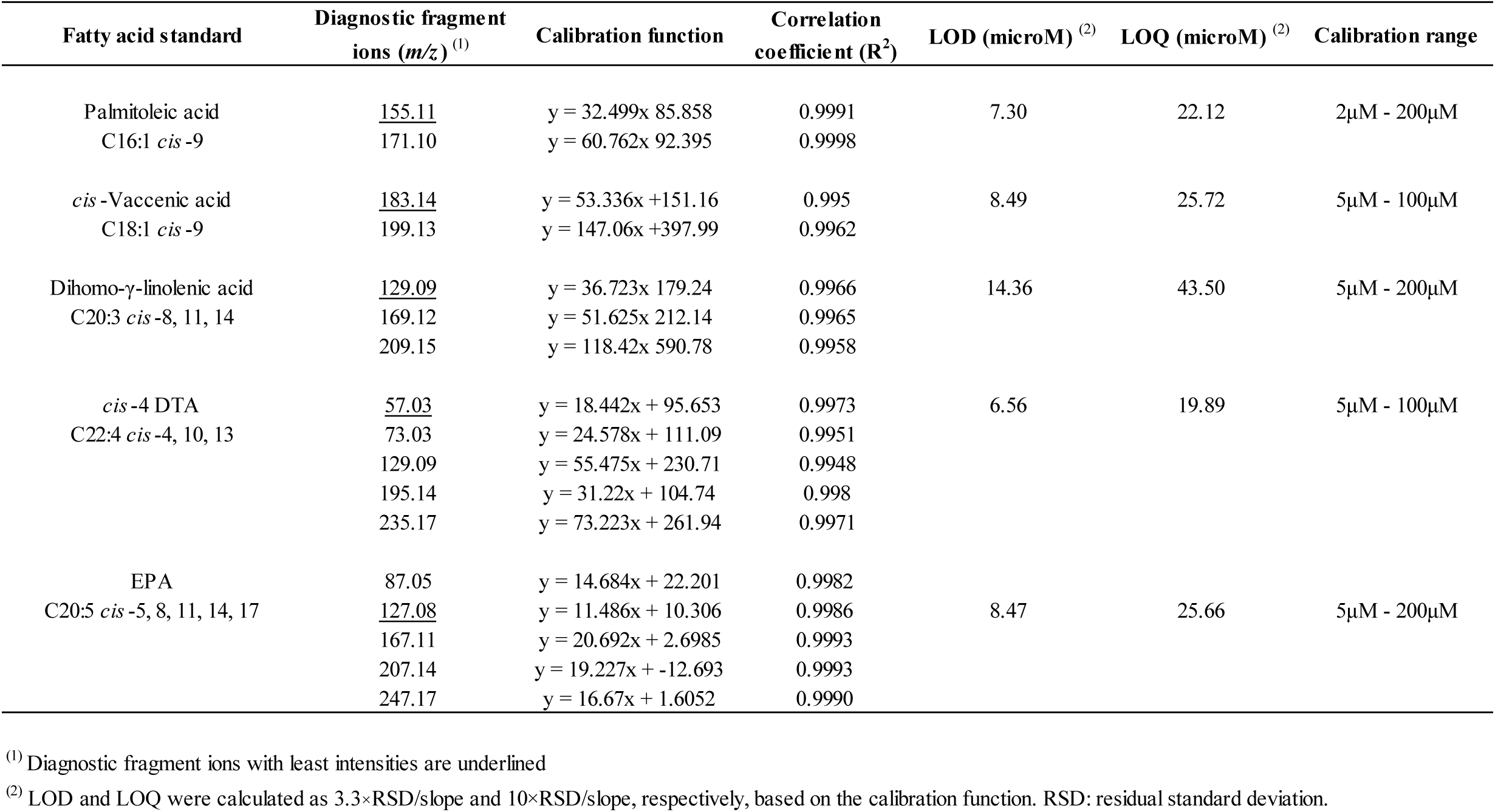
Analytical performances of MUFA and PUFA species.

We also evaluated the intraday (n = 5) and interday (n = 5) reproducibilities of diagnostic fragment ions using standard solutions of *cis*-vaccenic acid and γ-linolenic acid at two different concentrations (10 μM and 100 μM). The intraday precision, expressed as relative standard deviation (RSD), was less than 10% for most of the fragments (Table 2). The interday precision of RSD without internal standard (IS; ^13^C18:1 *cis*-9) was high; however, it improved upon normalization using IS (Table 2).

**Table 2.** Intraday and Interday precisions associated with this method.

| **Fatty acid standard** | **Concentration** | **Diagnostic fragments**<br>**(*m/z*)** | **Intraday RSD (%)**<br>**without I.S.** | **Intraday RSD (%)**<br>**with I.S.<sup>(1)</sup>** | **Interday RSD (%)**<br>**without I.S.** | **Interday RSD (%)**<br>**with I.S.<sup>(1)</sup>** |
| --- | --- | --- | --- | --- | --- | --- |
| *cis*-Vaccenic acid<br>C18:1 *cis*-9 | 10μM | 183.14 | 5.9% | 6.2% | 26.5% | 8.3% |
|  |  | 199.13 | 6.9% | 5.4% | 26.4% | 4.7% |
|  | 100μM | 183.14 | 4.2% | 7.3% | 28.0% | 9.2% |
|  |  | 199.13 | 3.6% | 8.0% | 28.4% | 5.6% |
| γ-Linolenic acid<br>C18:3 *cis*-6, 9, 12 | 10μM | 129.09 | 1.3% | 4.0% | 35.7% | 13.7% |
|  |  | 141.09 | 8.3% | 9.8% | 31.2% | 6.1% |
|  |  | 181.12 | 11.8% | 12.7% | 32.4% | 6.7% |
|  | 100μM | 129.09 | 5.7% | 10.9% | 31.6% | 15.0% |
|  |  | 141.09 | 3.9% | 9.0% | 30.3% | 12.4% |
|  |  | 181.12 | 4.7% | 10.4% | 30.1% | 12.6% |

<sup>(1)</sup> Fragment ions of the internal standard, <sup>13</sup>C-labeled oleic acid, was used for normalization.

Relative quantification was evaluated with mixtures of FA isomers of MUFA (oleic acid and *cis*-vaccenic acid), and PUFA (α-linolenic acid γ-linolenic acid), with several different ratios of the two isomers (Table 3). The relative quantity of the two isomers showed good linearity in the range from 0.1 to 2.0. The above data indicate that the relative abundance of FA isomers can be quantified using this method.

**Table 3.** Analytical performances in relative quantification.

| **First isomer** | **Second isomer** | **Corresponding fragments of**<br>**2<sup>nd</sup>/1<sup>st</sup> isomers (*m/z*)** | **Calibration function** | **R2** | **Ratio range**<br>**(2<sup>nd</sup>/1<sup>st</sup> isomer)** |
| --- | --- | --- | --- | --- | --- |
| Oleic acid<br>C18:1 *cis*-9<br>100μM | *cis*-Vaccenic acid | 183.14/155.11 | $y = 0.5744x + 0.3371$ | 0.9982 | 0.1 to 2.0 |
| | C18:1 *cis*-11 | 199.13/171.10 | $y = 0.7582x + 0.0724$ | 0.9954 | |
|  | 10μM to 200μM |  |  |  |  |
| α-Linolenic acid<br>C18:3 *cis*-9, 12, 15<br>100μM | γ-Linolenic acid | 129.09/87.05 | $y = 0.5368x + 0.0217$ | 0.9995 | 0.1 to 2.0 |
| | C18:3 *cis*-6, 9, 12 | 141.09/183.14 | $y = 0.4983x + 0.1484$ | 0.9876 | |
| | 10μM to 200μM | 181.12/223.17 | $y = 0.5328x + 0.0592$ | 0.9988 | |

### 3.5. Ionization and fragmentation mechanisms

The analytical mechanisms of this method were also studied. To elucidate the origin of oxygen in the epoxide and peroxide groups of the FAs formed by the plasma-ESI, we used ^18^O-containing water (H_2_^18^O) as a solvent. *Cis*-4 DTA was dissolved in a 1:1 solution of H_2_^18^O and acetonitrile, and was examined by direct infusion assay (additives like ammonium acetate and ammonia solution were not supplemented to eliminate any contaminant of regular water). Epoxide or peroxide formed by plasma-ESI had the same *m/z* values as those obtained using regular water (Figure 7A), indicating that the oxygen in epoxide or peroxide groups did not came from the solvent, but probably from the surrounding air as in the case of a previous report [15].

**Figure 7.**
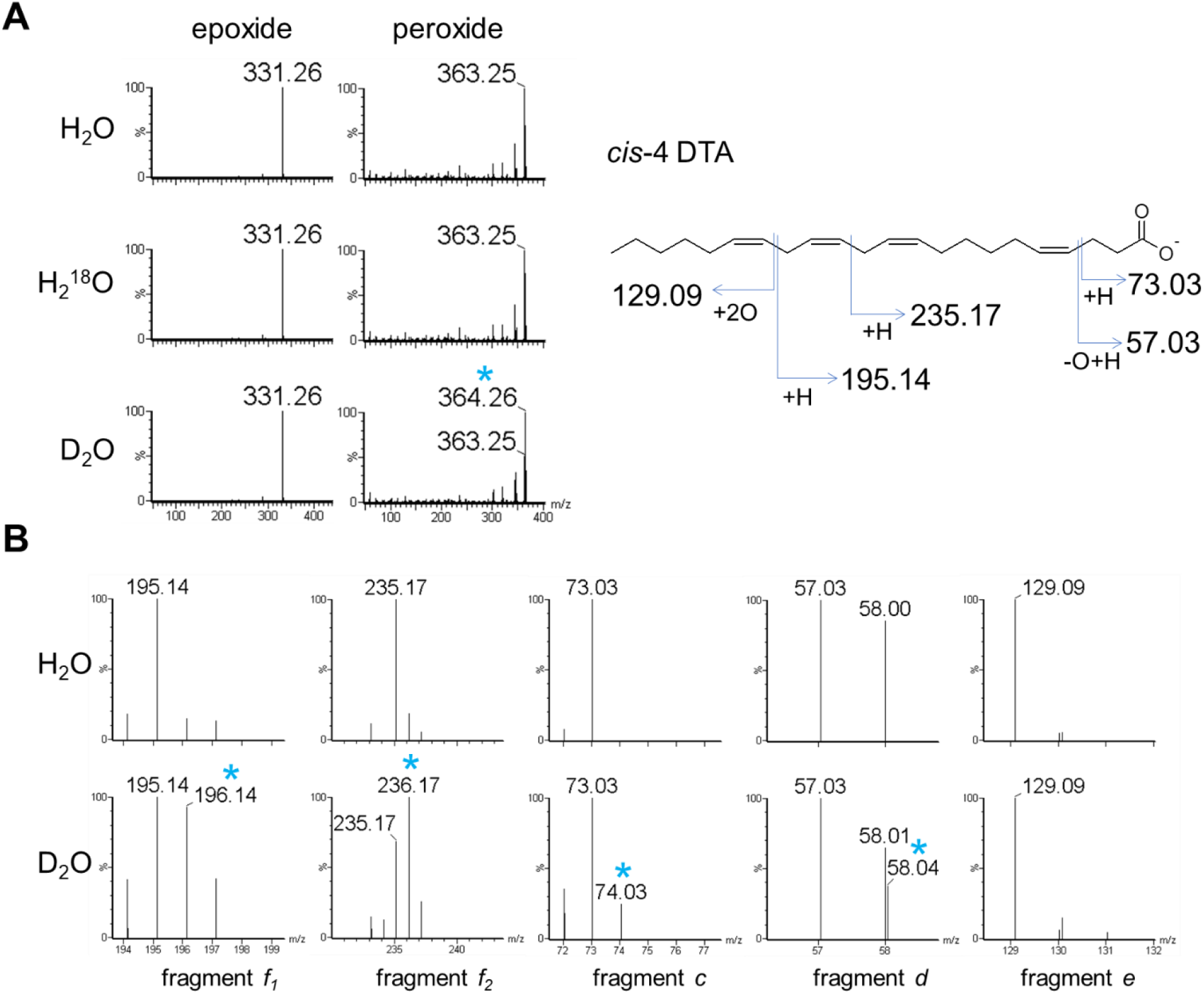
Mechanistic background of epoxide/peroxide formation and fragmentation. (A) Mass spectra of the epoxide and peroxide of *cis*-4 DTA. There is no mass increase of the epoxide produced with ^18^O-containing water (H_2_^18^O) or deuterium oxide (D_2_O), compared with regular water (H_2_O). There is no mass increase in the peroxide with H_2_^18^O; however, there is a mass increase by one unit with D_2_O (asterisk). (B) Diagnostic fragment ions of *cis*-4 DTA. The alpha fragments (*i.e.* m/z 195.14, 235.17, 73.03, and 57.03) show mass increase by one unit (asterisks) when D_2_O is used as the solvent, in contrast to the omega fragment that does not show mass increase.

We next used deuterium oxide (D_2_O) as a solvent to reveal the origin of the added hydrogen in the peroxide group and in some fragment ions (*i.e.* diagnostic fragment ions *c*, *d*, *e*, and *f*; see Figure 1C and D) by the same direct infusion assay. We found peroxidized *cis*-4 DTA and *cis*-vaccenic acid had increased *m/z* values by one unit, indicating that the solvent was the origin of the hydrogen in the peroxide group (Figure 7 and Figure S7, respectively). The diagnostic alpha fragment ions (diagnostic fragment ions *c*, *d*, and *f*) also had increased *m/z* values by one unit, again indicating that the hydrogen added to the fragments was derived from the solvent. On the other hand, the omega fragment (diagnostic fragment *e*) did not have increased *m/z* value. Taken together, these data indicate that the hydrogen in the peroxide group is included in the diagnostic fragment ions in the plasma-ESI method.

### 3.6. Analysis of unsaturated fatty acids from human fibroblasts

The developed method was subsequently applied to biological samples. FAs were extracted from human fibroblasts (NB1RGB) via acid hydrolysis and separated using a reverse-phase gradient LC (see Materials and Methods). The quadrupole was set to introduce epoxidized MUFA species (epoxi-MUFAs), including C16:1, C18:1, C20:1, C22:1, C24:1, and C26:1, in separate time windows. The introduced epoxi-MUFAs were subsequently fragmented by CID. The entire ions ranging from *m/z* 100 to 1000 were also scanned and recorded simultaneously in another channel to detect the non-epoxidized FAs. As reported previously [24], multiple peaks can be observed for some of the MUFAs in their mass chromatograms (Figure 8A; Figure S8A, S9A), which may represent FA isomers with different double bond positions. This observation was confirmed by characterizing the isomers in each peak. In the mass chromatogram of C16:1, two peaks were observed with incomplete base separation. Upon CID fragmentation, three C16:1 isomers (Figure 8) were observed. In the first peak, *ω*-7 (*cis*-9) isomer yielding fragment ions at *m/z* 155.11 and 171.10 was detected. At the beginning area of the second peak, *ω*-9 (*cis*-7) isomer was observed yielding fragments at *m/z* 143.07 and 127.08, while *ω*-10 (*cis*-6) isomer yielding fragments at *m/z* 113.06 and 129.06 was found throughout most of the second peak. The retention order of the isomers followed previous observations where the FAs with more proximal double bonds to the omega carbon eluted faster than those with more distal double bonds.

**Figure 8.**
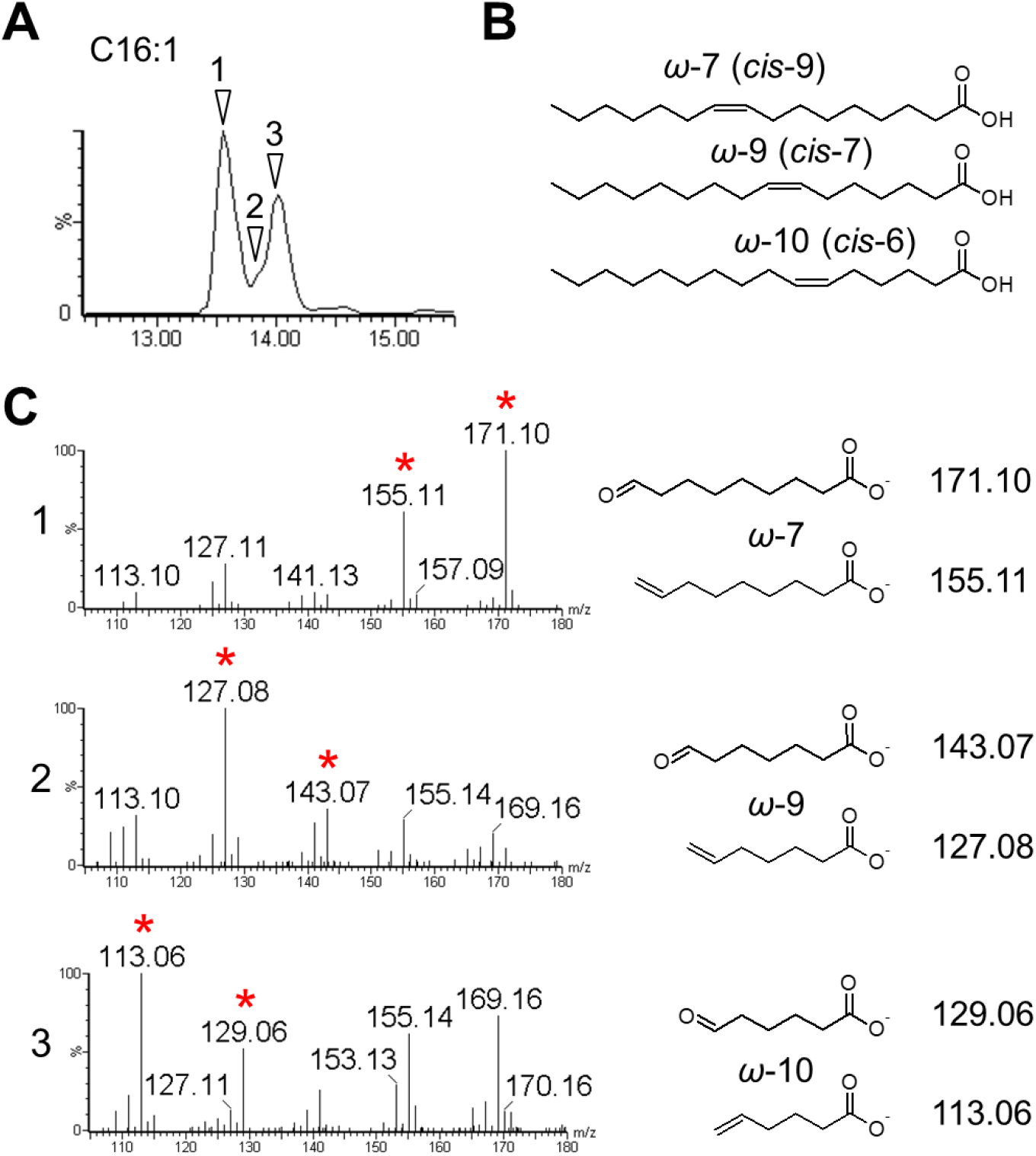
Identification of multiple C16:1 FA isomers in human fibroblasts. (A) Mass chromatogram of C16:1 (*m/z* 253.22). The time points where each C16:1 isomer was identified are indicated by arrowheads. (B) Detected C16:1 isomers from the analysis. Three C16:1 isomers with double bonds at *ω*-7, *ω*-9, and *ω*-10 were discovered. (C) Tandem mass spectra and diagnostic fragment structure of each C16:1 isomer at the indicated time points in (A) are shown and diagnostic fragment ions are indicated by asterisks.

From the C18:1 assay, four isomers were detected (Figure 9), with the most abundant being oleic acid (C18:1 *ω*-9, *cis*-9), which yielded a fragment ion pair at *m/z* 171.10 and 155.11. The other C18:1 isomers were *ω*-7 (*cis*-11) (producing a fragment ion pair at *m/z* 199.13 and 183.14), *ω*-10 (*cis*-8) (*m/z* 157.09 and 141.09), and *ω*-12 (*cis*-6) (*m/z* 129.05 and 113.06). *ω*-7, *ω*-9, and *ω*-10 isomers were observed in the first major peak, while *ω*-12 isomer was detected in the minor second peak (Figure 9). Comparing the retention times of the isomers, the elution order from the earliest to the latest was *ω*-7, *ω*-9, *ω*-10, and *ω*-12 isomers (Figure 9). Even though four isomers of C18:1 were detected, the high abundance of oleic acid (*ω*-9 isomer) embedded the other isomers within a single large peak, highlighting the importance of isomer segregation. Similarly, multiple isomers were observed in the assays of C20:1 to C26:1. The isomers with their diagnostic fragment ions are shown in Figures S8–S11. The double bond position of MUFAs from biological samples, thus, were confirmed by this method.

**Figure 9.**
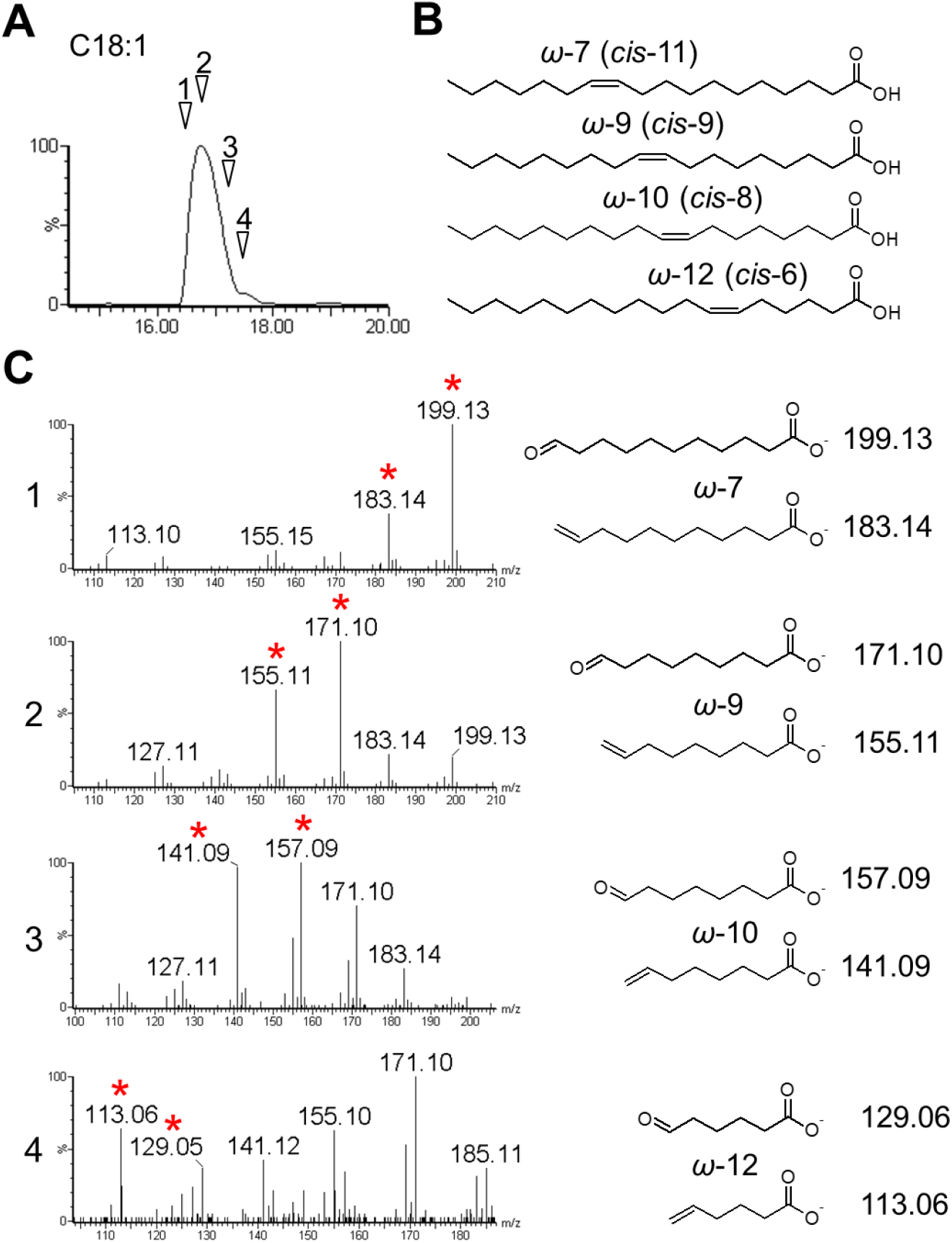
Identification of multiple C18:1 isomers from human fibroblasts. The same FA samples shown in Figure 8 were analyzed. (A) Mass chromatogram of C18:1 (*m/z* 281.25). Analytical time points where each isomer was detected are indicated. (B) A total of four C18:1 isomers each possessing double bonds at *ω*-7, *ω*-9, *ω*-10, and *ω*-12 were discovered. (C) Tandem mass spectra and structures of the diagnostic ions at the indicated time points in (A). The diagnostic fragment ions are indicated by asterisks.

Finally, PUFA isomers from a clinical sample were analyzed. FAs were extracted from fibroblasts of a human patient with Zellweger syndrome (ZS) and from control fibroblasts (F-12 and NB1RGB, respectively) [12]. In ZS, the biosynthesis of peroxisomes is impaired (Figure 10A) and the FA composition fluctuates due to metabolic failures in peroxisomes. In our previous study, multiple peaks were obtained for C20:3 FA upon LC separation, where each peak was suggested to be a distinct C20:3 isomer. Moreover, the isomeric abundance differed between ZS and control fibroblasts, although the exact double bond position remained undetermined [24]. By analyzing the FAs from the fibroblast samples, multiple peaks for C20:3 were obtained (Figure 10B). Peroxidation of C20:3 followed by CID fragmentation indicated that the first peak included a C20:3 isomer with the double bonds at *cis*-8, 11, and 14 positions in ZS and control fibroblasts (Figure 10B-D). Moreover, an additional isomer with double bonds at *cis*-7, 10, and 13 positions was found in both samples but reduced in ZS fibroblasts (Figure 10C and 10D, Figure S12). The second peak corresponded to only a single isomer with double bonds at *cis*-5, 8, and 11 positions found in both samples but again reduced in ZS fibroblasts (Figure 10C and 10D, Figure S12). These results confirmed our previous observation of control and ZS fibroblasts [24]. These data indicate that the PUFA isomers in biological samples can be distinguished. This is the first example that showed differential abundances of specific PUFA isomers in ZS samples, highlighting the usefulness of the method reported herein for medical applications.

**Figure 10.**
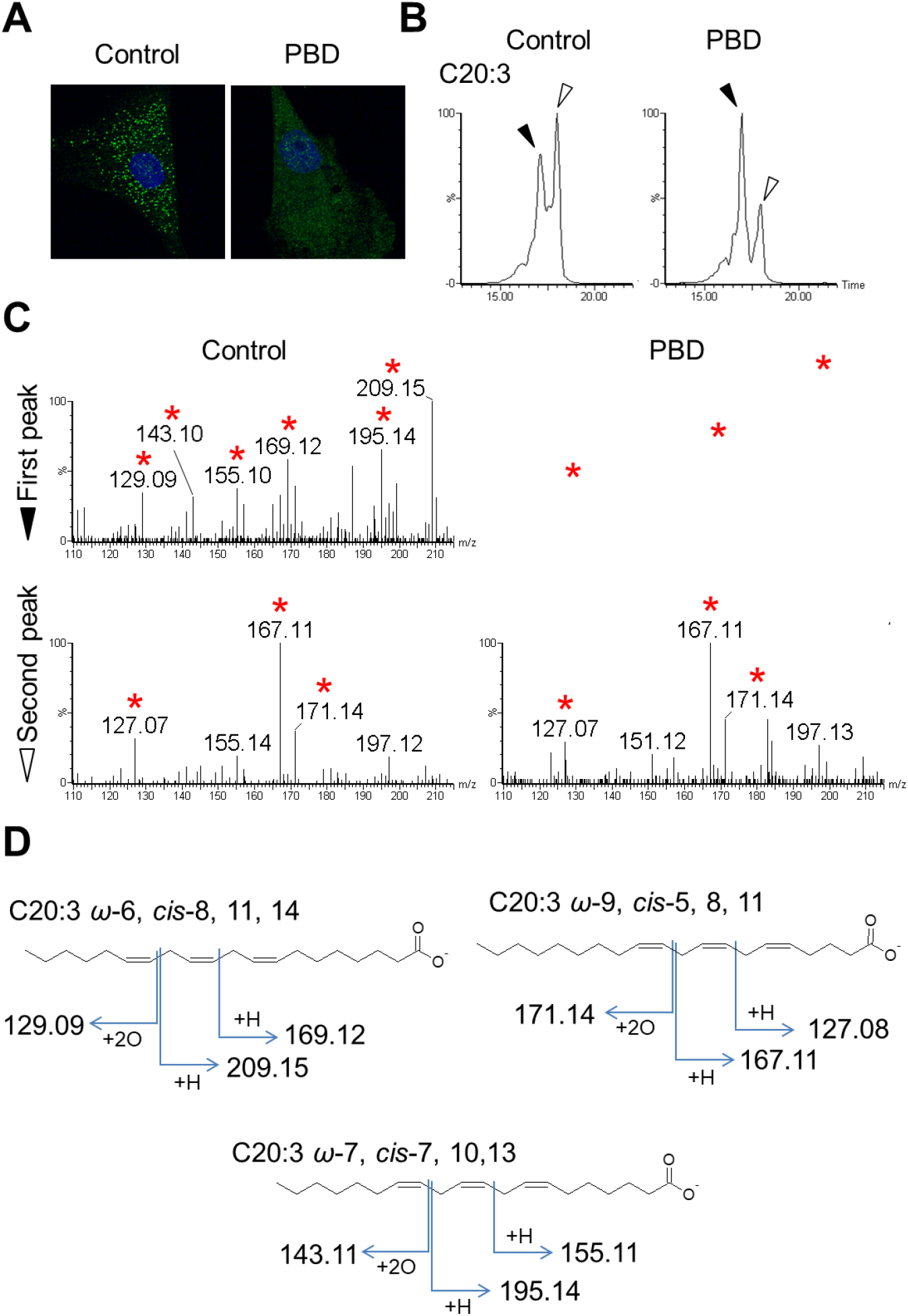
Identification and comparison of the C20:3 PUFA species between control fibroblasts and fibroblasts from a ZS patient. (A) Anti-catalase staining (green) of the fibroblasts. The nuclei were stained with DAPI in blue. Vesicular peroxisomes are visible only in the control fibroblast. (B) Mass chromatogram of C20:3 (*m/z* 337.24) where the peak tops are indicated by black (first peak) and white (second peak) arrowheads. (C) Tandem mass spectra at each peak where the diagnostic fragment spectra are indicated by asterisks. (D) Chemical structures and diagnostic fragmentation patterns of the detected C20:3 species are shown.

## 4. CONCLUSIONS

Determination of the positions and number of double bonds, as well as the chain length, is crucial for understanding the precise physiological functions of each FA species. Many methods have been established for the positional determination of double bonds by modifying the target FAs at the double bond positions or the terminal carboxyl group to yield diagnostic fragment ions [8–11,13–23]. CID-mediated fragmentation on ESI-MS or EI-mediated fragmentation on GC-MS of unsaturated FAs cannot always produce informative product ions to determine the double bond positions, especially in case of PUFAs with many double bonds. Moreover, the derivatization of FAs affects their chromatographic retention time, compromising the interpretation of chromatographic results. Herein, a method of post-column epoxidation and peroxidation of unsaturated FAs using corona discharge plasmatization of chromatographic solvent is reported. This method is advantageous for samples containing mixed FAs, such as biological samples. The post-column epoxidation/peroxidation does not compromise the LC, so the chain length and number of double bonds in each FA in the mixture can be easily determined from the LC results [24]. The drawback of this method, presently, is its relatively low sensitivity, attributable to the poor epoxidation/peroxidation efficiency and target ionization. Because a pre-equipped setup for conventional ESI was used in this study, the development of a specialized setup may improve the sensitivity for detecting FA species with low abundance. Alternatively, addition of an epoxidizing/peroxidizing agent to the solvent would improve the sensitivity of this method. In addition, post-column modification of target molecules with solvent plasmatization can be applied to other types of molecules for structural analysis. In conclusion, the method reported herein enabled a thorough characterization of FA species, *i.e.* chain length, number of double bonds, and position of double bonds, and has the potential to greatly impact medical, biological, and agricultural sciences as well as the food industry.

## Abbreviations

FA: fatty acid
MUFA: mono-unsaturated fatty acid
PUFA: poly-unsaturated fatty acid
VLCFA: very-long-chain fatty acid
ZS: Zellweger syndrome

## Author contributions

ST and NS designed research, ST and KT performed experiments, TY contributed on theoretical analysis, and NS prepared the biopsy sample.

## Acknowledgements

This work was supported by research aid funds from the Koshiyama Foundation for Promotion of Science and Technology, and the Takahashi Industrial and Economic Research Foundation.

## References

[1] Food and Agriculture Organization of the United Nations, Fats an fatty acids in human nutrition - Report of an expert consultation, 2010.

[2] C. Ferreri, A. Masi, A. Sansone, G. Giacometti, A. Larocca, G. Menounou, R. Scanferlato, S. Tortorella, D. Rota, M. Conti, S. Deplano, M. Louka, A. Maranini, A. Salati, V. Sunda, C. Chatgilialoglu, Fatty Acids in Membranes as Homeostatic, Metabolic and Nutritional Biomarkers: Recent Advancements in Analytics and Diagnostics, Diagnostics. 7 (2016) 1. https://doi.org/10.3390/diagnostics7010001.

[3] de Carvalho, M.J. Caramujo, The various roles of fatty acids, Molecules. 23 (2018). https://doi.org/10.3390/molecules23102583.

[4] Y. Isobe, M. Arita, Identification of novel omega-3 fatty acid-derived bioactive metabolites based on a targeted lipidomics approach, J. Clin. Biochem. Nutr. 55 (2014) 79–84. https://doi.org/10.3164/jcbn.14-18.

[5] J. Miyata, M. Arita, Role of omega-3 fatty acids and their metabolites in asthma and allergic diseases, Allergol. Int. 64 (2015) 27–34. https://doi.org/10.1016/j.alit.2014.08.003.

[6] M.P. Wymann, R. Schneiter, Lipid signalling in disease., Nat. Rev. Mol. Cell Biol. 9 (2008) 162–76. https://doi.org/10.1038/nrm2335.

[7] J.J. Liu, P. Green, J. John Mann, S.I. Rapoport, M.E. Sublette, Pathways of polyunsaturated fatty acid utilization: Implications for brain function in neuropsychiatric health and disease, Brain Res. 1597 (2015) 220–246. https://doi.org/10.1016/J.BRAINRES.2014.11.059.

[8] H. R. Buser, H. Arn, P. Guerin, S. Rauscher, Determination of double bond position in mono-unsaturated acetates by mass spectrometry of dimethyl disulfide adducts, Anal. Chem. 55 (1983) 818–822. https://doi.org/10.1021/ac00257a003.

[9] M. Vincent, G. Guglielmetti, G. Cassani, C. Tonini, Determination of double-bond position in diunsaturated compounds by mass spectrometry of dimethyl disulfide derivatives, Anal. Chem. 59 (1987) 694–699. https://doi.org/10.1021/ac00132a003.

[10] K. Yamamoto, A. Shibahara, T. Nakayama, G. Kajimoto, Determination of double-bond positions in methylene-interrupted dienoic fatty acids by GC-MS as their dimethyl disulfide adducts, Chem. Phys. Lipids. 60 (1991) 39–50. https://doi.org/10.1016/0009-3084(91)90013-2.

[11] A. Shibahara, K. Yamamoto, A. Kinoshita, B.L. Anderson, An Improved Method for Preparing Dimethyl Disulfide Adducts for GC/MS Analysis, J. Am. Oil Chem. Soc. 85 (2008) 93–94. https://doi.org/10.1007/s11746-007-1169-7.

[12] S. Takashima, K. Toyoshi, T. Itoh, N. Kajiwara, A. Honda, A. Ohba, S. Takemoto, S. Yoshida, N. Shimozawa, Detection of unusual very-long-chain fatty acid and ether lipid derivatives in the fibroblasts and plasma of patients with peroxisomal diseases using liquid chromatography-mass spectrometry, Mol. Genet. Metab. 120 (2017) 255–268. https://doi.org/10.1016/j.ymgme.2016.12.013.

[13] V. Vrkoslav, M. Háková,K. Pecková, K. Urbanová,J. Cvačka, Localization of Double Bonds in Wax Esters by High-Performance Liquid Chromatography/Atmospheric Pressure Chemical Ionization Mass Spectrometry Utilizing the Fragmentation of Acetonitrile-Related Adducts, Anal. Chem. 83 (2011) 2978–2986. https://doi.org/10.1021/ac1030682.

[14] V. Vrkoslav, J. Cvačka, Identification of the double-bond position in fatty acid methyl esters by liquid chromatography/atmospheric pressure chemical ionisation mass spectrometry, J. Chromatogr. A. 1259 (2012) 244–250. https://doi.org/10.1016/J.CHROMA.2012.04.055.

[15] Y. Zhao, H. Zhao, X. Zhao, J. Jia, Q. Ma, S. Zhang, X. Zhang, H. Chiba, S.P. Hui, X. Ma, Identification and Quantitation of C=C Location Isomers of Unsaturated Fatty Acids by Epoxidation Reaction and Tandem Mass Spectrometry, Anal. Chem. 89 (2017) 10270–10278. https://doi.org/10.1021/acs.analchem.7b01870.

[16] W. Cao, X. Ma, Z. Li, X. Zhou, Z. Ouyang, Locating Carbon–Carbon Double Bonds in Unsaturated Phospholipids by Epoxidation Reaction and Tandem Mass Spectrometry, Anal. Chem. 90 (2018) 10286–10292. https://doi.org/10.1021/acs.analchem.8b02021.

[17] L. Wan, G. Gong, H. Liang, G. Huang, In situ analysis of unsaturated fatty acids in human serum by negative-ion paper spray mass spectrometry, Anal. Chim. Acta. 1075 (2019) 120–127. https://doi.org/10.1016/j.aca.2019.05.055.

[18] X. Ma, L. Chong, R. Tian, R. Shi, T.Y. Hu, Z. Ouyang, Y. Xia, Identification and quantitation of lipid C=C location isomers: A shotgun lipidomics approach enabled by photochemical reaction., Proc. Natl. Acad. Sci. U. S. A. 113 (2016) 2573–8. https://doi.org/10.1073/pnas.1523356113.

[19] R.C. Murphy, T. Okuno, C.A. Johnson, R.M. Barkley, Determination of Double Bond Positions in Polyunsaturated Fatty Acids Using the Photochemical Paternò-Büchi Reaction with Acetone and Tandem Mass Spectrometry, Anal. Chem. 89 (2017) 8545–8553. https://doi.org/10.1021/acs.analchem.7b02375.

[20] F. Tang, C. Guo, X. Ma, J. Zhang, Y. Su, R. Tian, R. Shi, Y. Xia, X. Wang, Z. Ouyang, Rapid In Situ Profiling of Lipid C=C Location Isomers in Tissue Using Ambient Mass Spectrometry with Photochemical Reactions, Anal. Chem. 90 (2018) 5612–5619. https://doi.org/10.1021/acs.analchem.7b04675.

[21] X. Xie, Y. Xia, Analysis of Conjugated Fatty Acid Isomers by the Paternò-Büchi Reaction and Trapped Ion Mobility Mass Spectrometry, Anal. Chem. 91 (2019) 7173–7180. https://doi.org/10.1021/acs.analchem.9b00374.

[22] W.C. Yang, J. Adamec, F.E. Regnier, Enhancement of the LC/MS analysis of fatty acids through derivatization and stable isotope coding, Anal. Chem. 79 (2007) 5150–5157. https://doi.org/10.1021/ac070311t.

[23] M. Wang, R.H. Han, X. Han, Fatty Acidomics: Global Analysis of Lipid Species Containing a Carboxyl Group with a Charge-Remote Fragmentation-Assisted Approach, Anal. Chem. 85 (2013) 9312–9320. https://doi.org/10.1021/ac402078p.

[24] S. Takashima, K. Toyoshi, N. Shimozawa, Analyses of the fatty acid separation principle using liquid chromatography mass spectrometry, Med. Mass Spectrom. 2 (2018) 1–13. https://doi.org/10.24508/mms.2018.06.002.

