## Supplemental Figure S1 to S12 for "A method to determine the double bond position in unsaturated fatty acids by solvent plasmatization using liquid chromatography-mass spectrometry"

**Supplemental data**

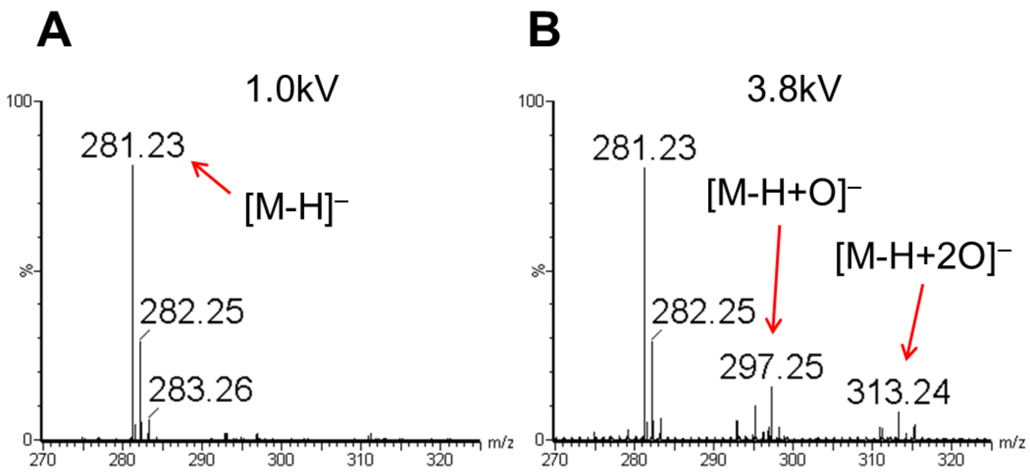


**Figure S1.** Plasma-facilitated modification of oleic acid in direct infusion assay. (A, B) Mass chromatogram of oleic acid with different capillary voltage. At l.0 kv (A), oleic acid was detected as deprotonated monovalent anion ($[M\text{-}H]^-$; calculated *m/z* of 281.25). At high voltage (*e.g.* 3.8 kv, B), epoxidized, deprotonated anion ($[M\text{-}H\text{+}O]^-$, *m/z* 297.24) and peroxidized, deprotonated anion ($[M\text{-}H\text{+}2O]^-$, *m/z* 313.24) were detected simultaneously.

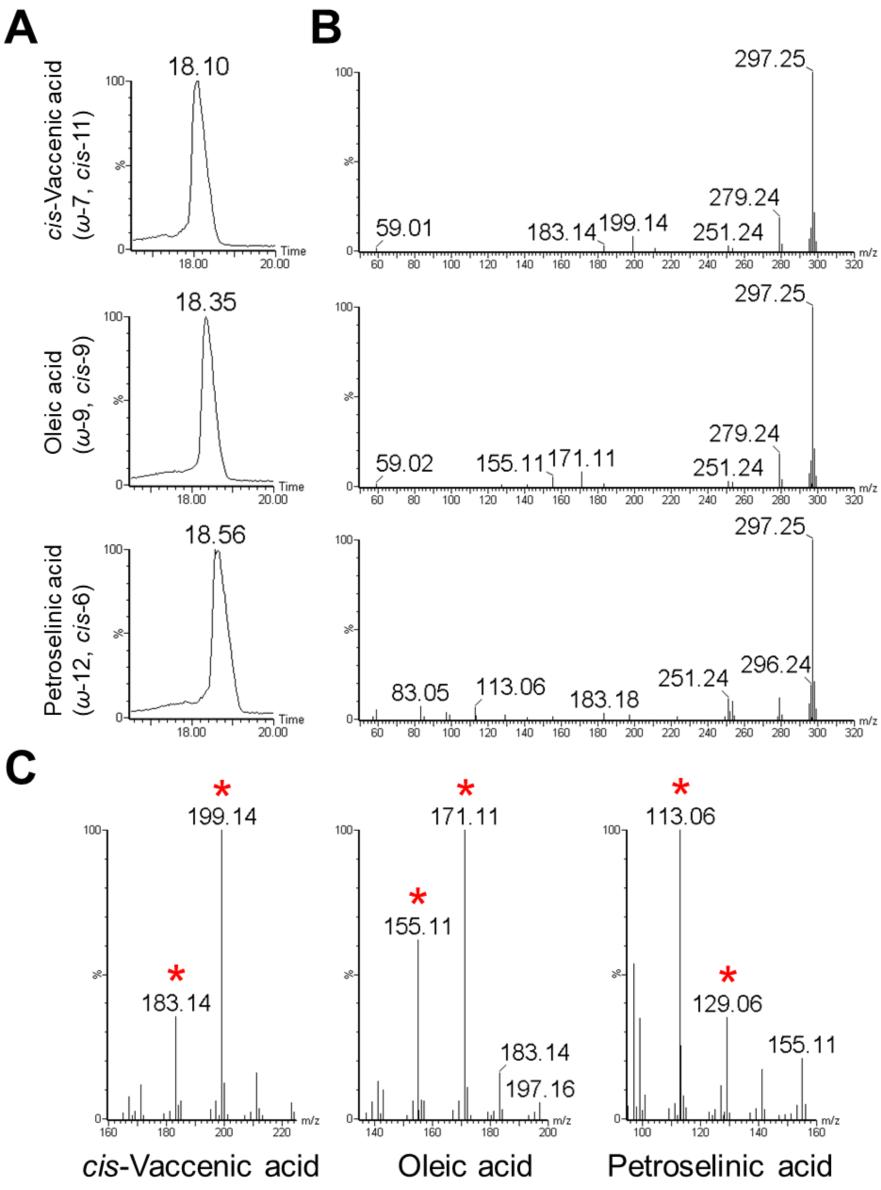


**Figure S2.** Structure determination of C18:1 FAs by LC-plasma-ESI-MS. (A) Mass chromatogram of oleic acid, *cis*-vaccenic acid, and petroselinic acid (*m/z* 281.25). Retention times are provided at the peak top. (B, C) Fragment spectra of the three FAs show the fragmentation pattern similar to the direct infusion assay (Figure 2). Diagnostic ions are indicated by asterisks in (C).

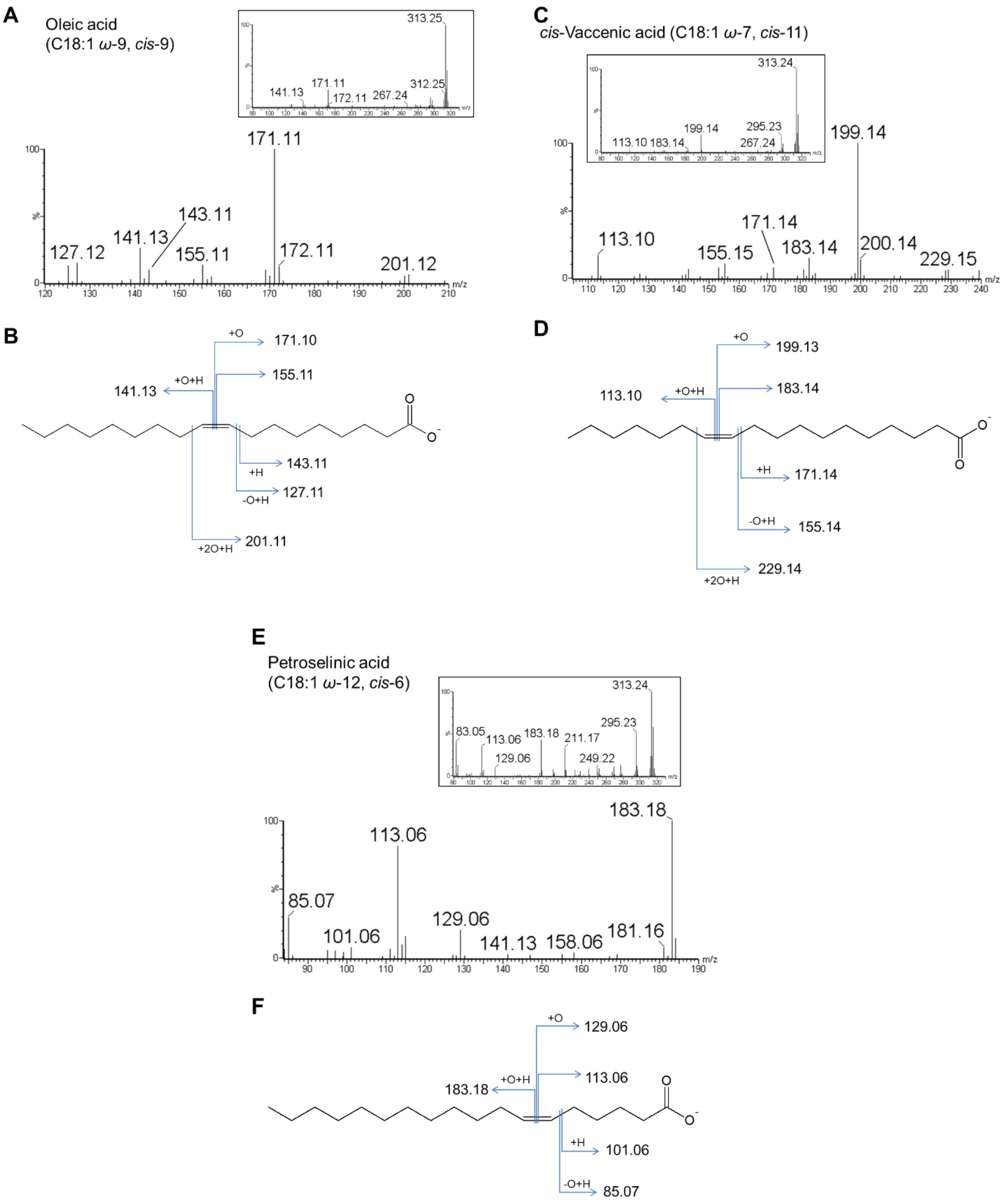


**Figure S3.** Fragmentation of C18:1 FAs. (A–F) Detailed fragment patterns of the peroxidized C18:1 FA species (*m/z* 313.24), such as oleic acid (A, B), *cis*-vaccenic acid (C, D), and petroselinic acid (E, F).

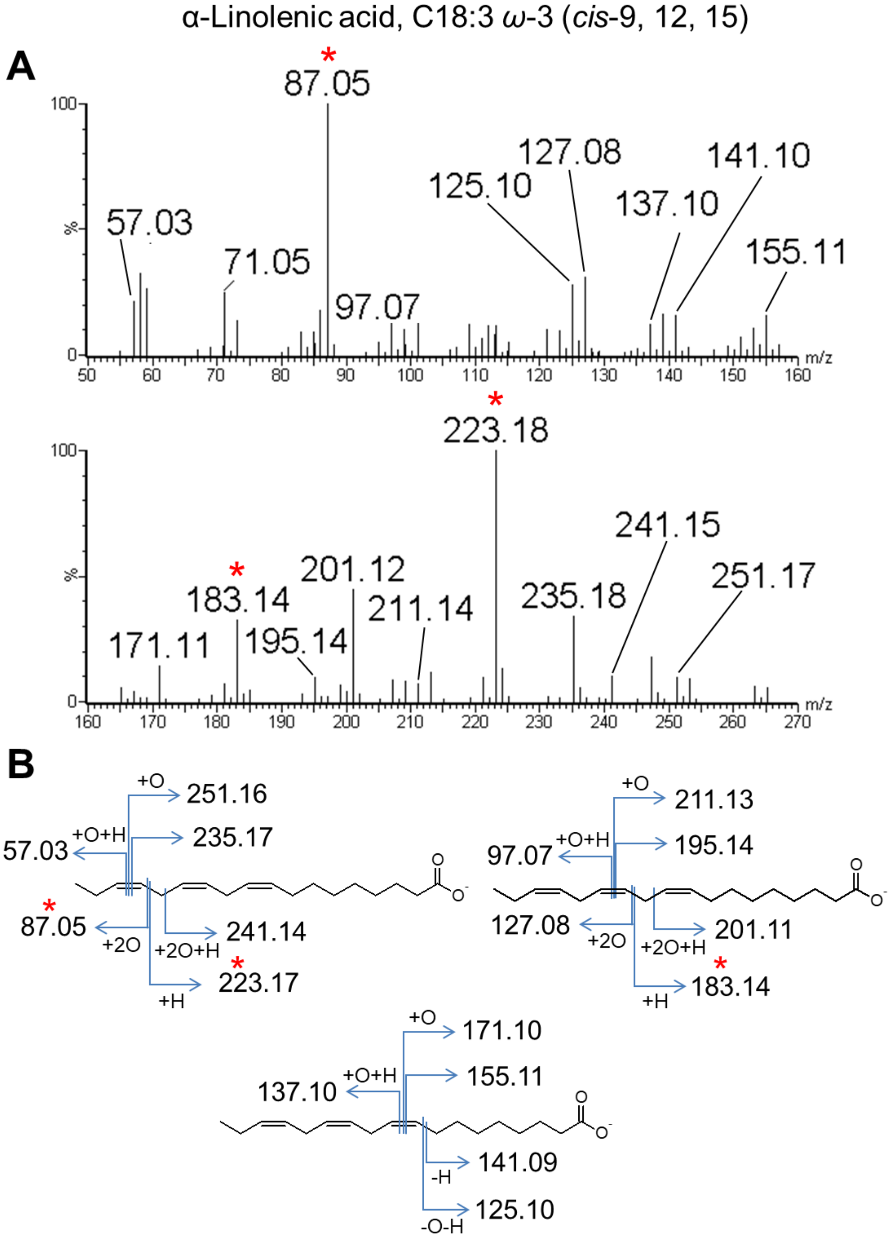


**Figure S4.** Detailed fragmentation pattern of α-linolenic acid. (A) Tandem mass spectrum of peroxidized α-linolenic acid. (B) Fragmentation patterns associated with each double bond are shown. Diagnostic fragment ions are marked by asterisks.

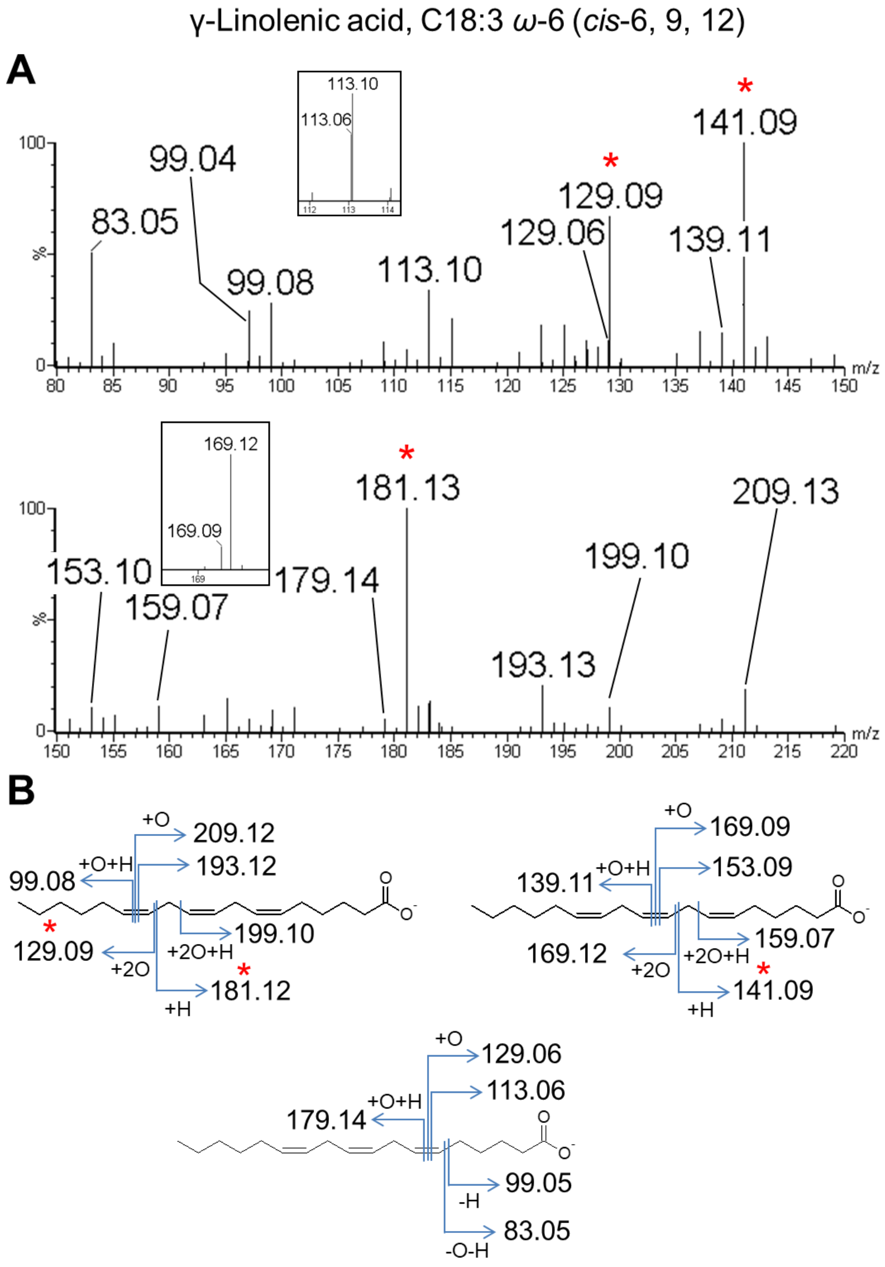


**Figure S5.** Detailed fragmentation pattern of γ-linolenic acid. (A) Tandem mass spectrum of peroxidized γ-linolenic acid. (B) Fragmentation patterns associated with each double bond are shown. Diagnostic fragment ions are marked by asterisks.

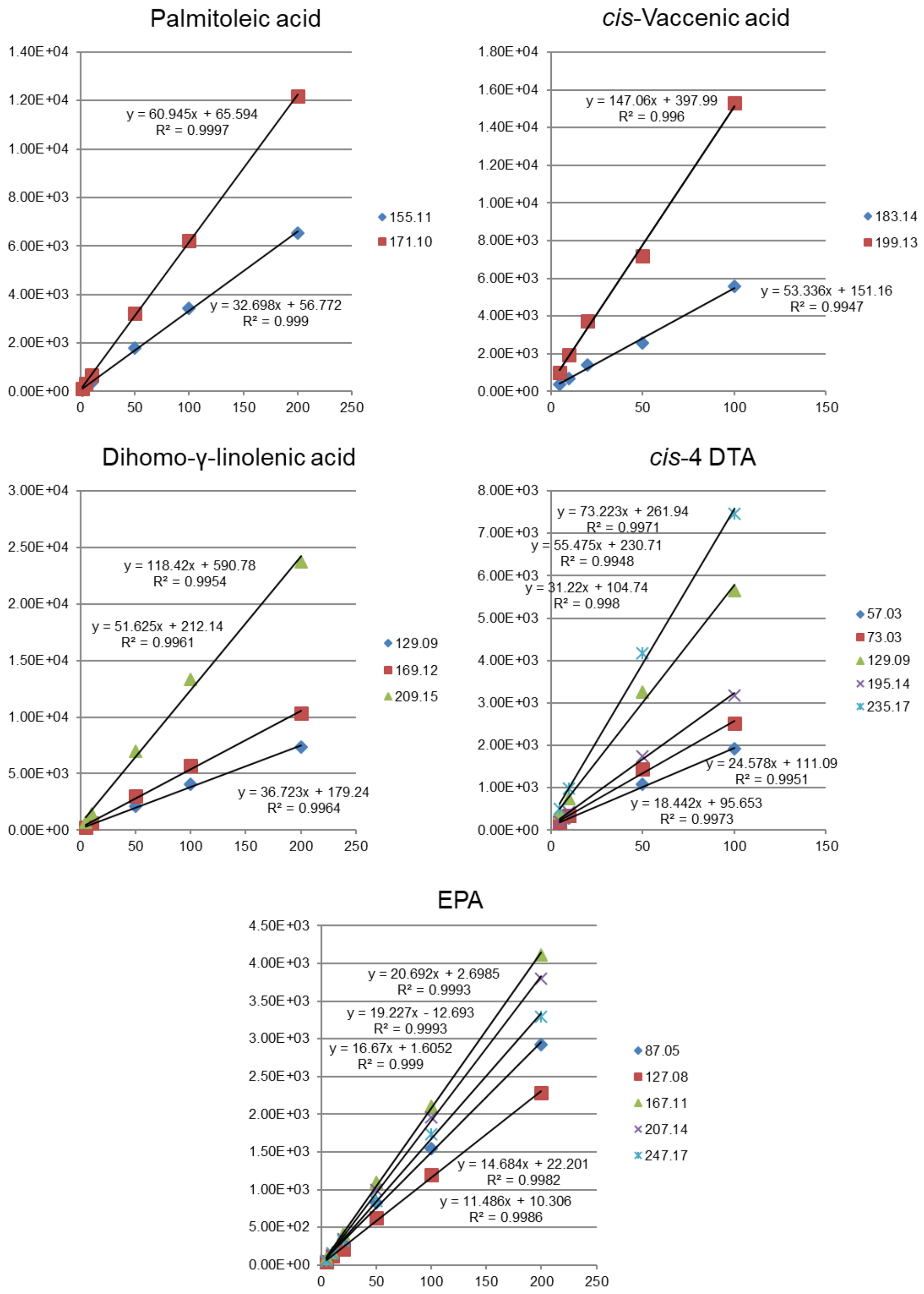


**Figure S6.** Calibration curves for several FA species. Diagnostic fragment ions of MUFA, such as palmitoleic acid (C16:1 *cis*-9) and *cis*-vaccenic acid (C18:1 *cis*-11), as well as PUFA, dihomo-γ-linolenic acid (C20:3 *cis*-8, 11, 14), *cis*-4 DTA (C22:4 *cis*-4, 10, 13, 16), and EPA (C20:5 *cis*-5, 8, 11, 14, 17), were plotted individually. Calibration function and correlation coefficient ($R^2$) are shown. The *x*-and *y*-axes represent FA concentration and intensity, respectively.

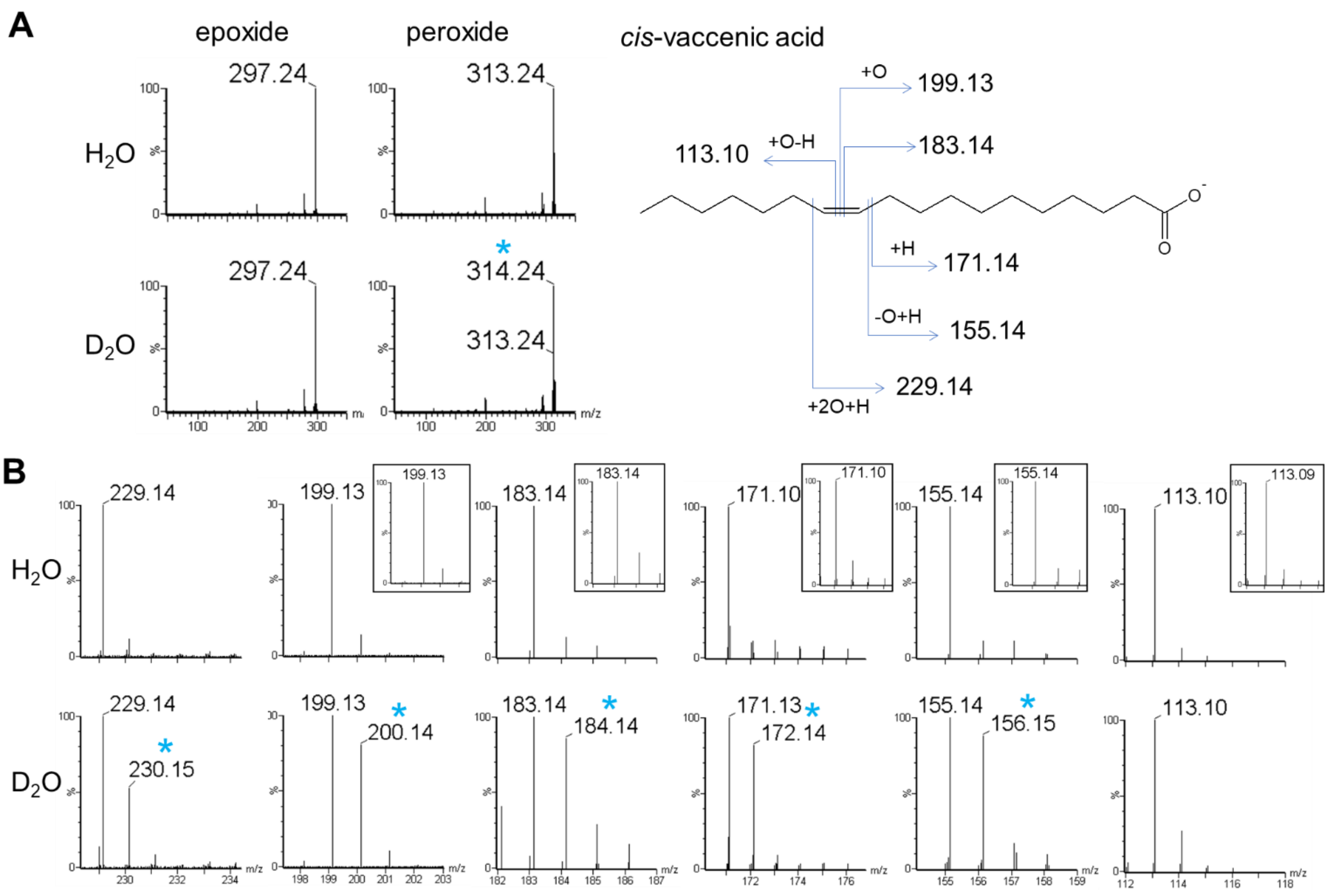


**Figure S7.** Fragmentation of *cis*-vaccenic acid with $D_2O$. (A) Mass spectra of epoxide and peroxide of *cis*-vaccenic acid are shown. There is no increase in the mass of the epoxide produced with $D_2O$ as a solvent or regular water ($H_2O$). There is a one-unit increase in the mass of the peroxide produced with $D_2O$ (asterisk). (B) Fragment ions of the peroxide of *cis*-vaccenic acid with $D_2O$ and $H_2O$. The alpha fragment ions (*m/z* 229.14, 199.13, 183.14, 171.14, and 155.14) show mass increase by one unit (asterisks) when $D_2O$ is used as a solvent, in contrast to the omega fragment (*m/z* 113.10) that does not show clear mass increase. The mass increase of the alpha fragment ions is peroxide-specific, such that the equivalent fragments of epoxide with $D_2O$ (insets) do not show mass increase (the fragment ion with *m/z* 229.14 was not produced from epoxide).

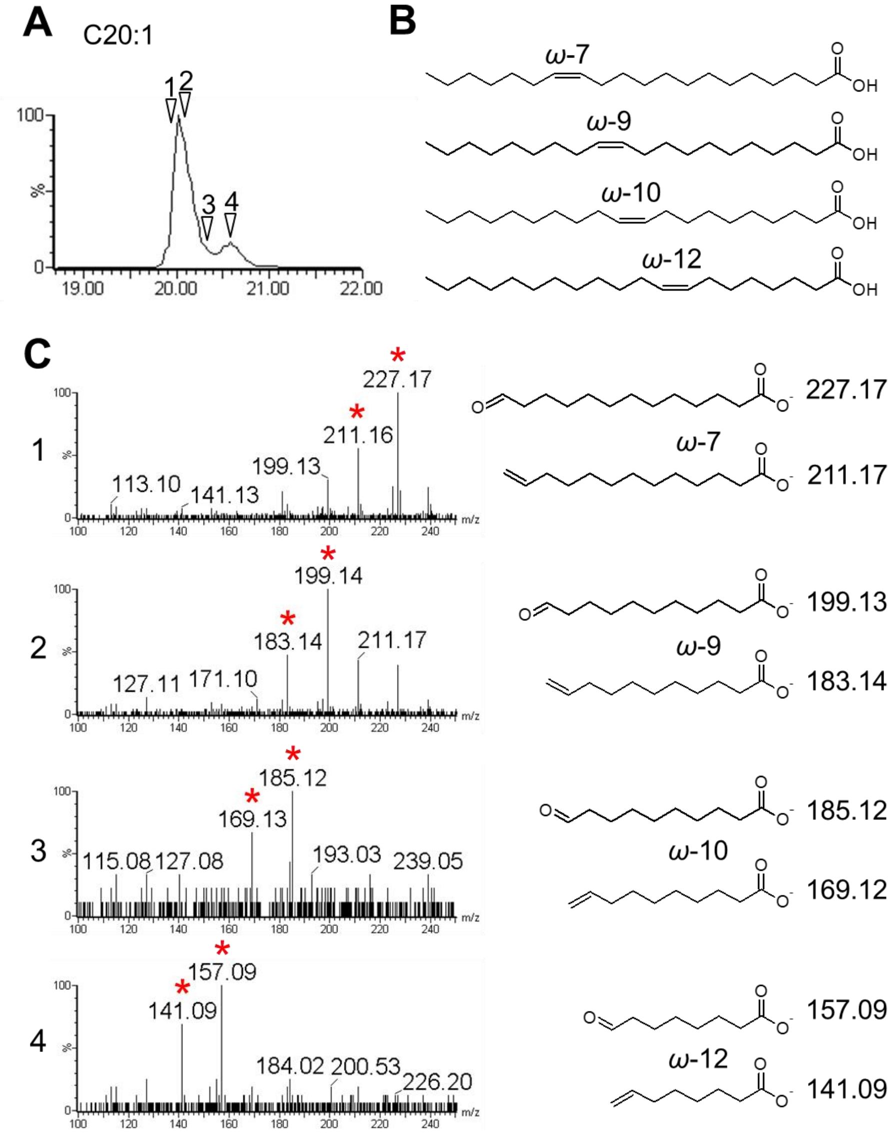


**Figure S8.** Identification of multiple C20:1 FA isomers from human fibroblasts. The same FA sample as shown in Figure 7 was analyzed. (A) Mass chromatogram of C20:1 on LC-ESI-MS (*m/z* 309.28). Analytical time points are indicated by arrowheads. (B) Four C20:1 isomers with double bonds either at *ω*-7, *ω*-9, *ω*-10, or *ω*-12 were detected. (C) Tandem mass spectra and structures of the diagnostic ions at the indicated time points in (A) are shown. The diagnostic fragment ions are indicated by asterisks in the spectral graph.

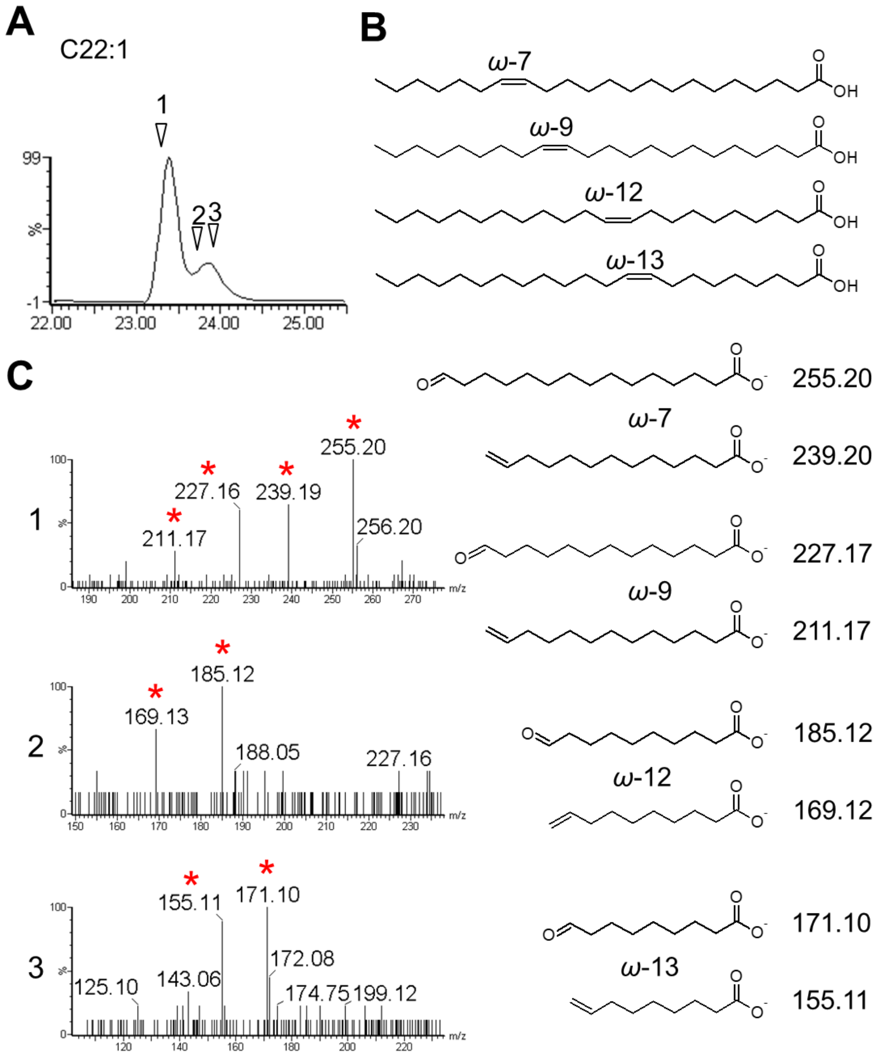


**Figure S9.** Identification of multiple C22:1 FA isomers from human fibroblasts. The same FA sample as shown in Figure 7 was analyzed. (A) Mass chromatogram of C22:1 on LC-ESI-MS (*m/z* 337.31). Analytical time points are indicated. (B) Four C22:1 isomers with double bond either at *ω*-7, *ω*-9, *ω*-12, or *ω*-13 position were detected. (C) Tandem mass spectra and structures of the diagnostic ions at the indicated time points in (A) are shown. Diagnostic fragment ions are indicated by asterisks in the spectral graph.

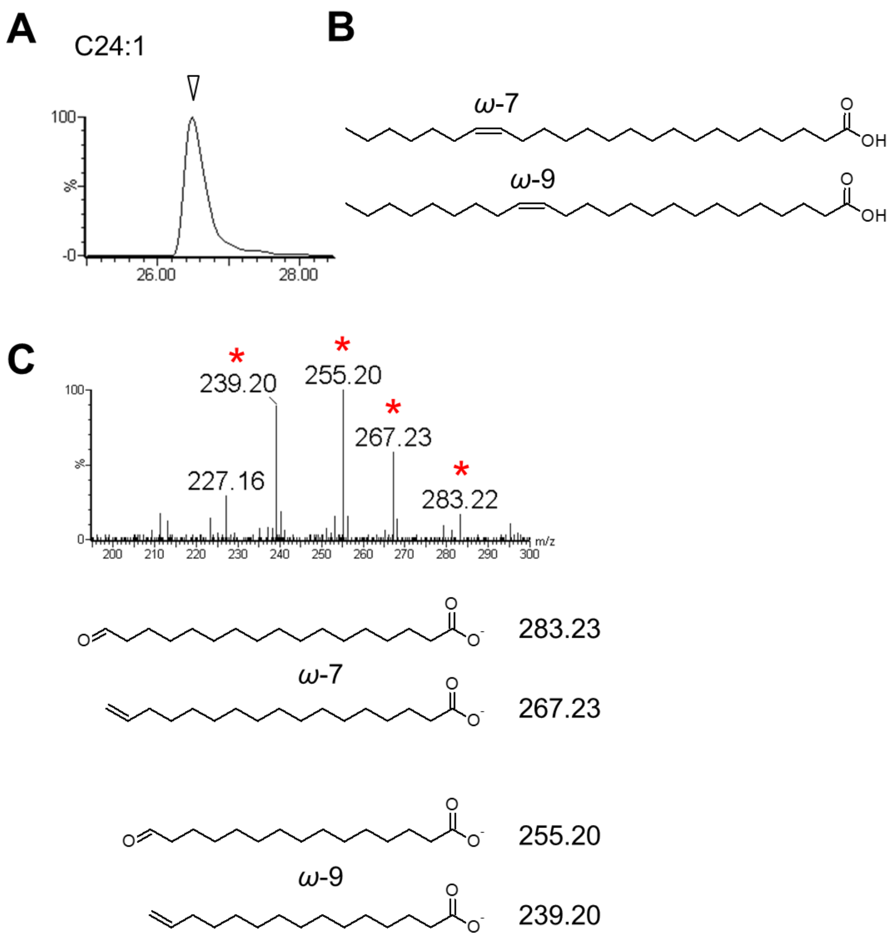


**Figure S10.** Identification of C24:1 FA isomers from human fibroblasts. The same FA sample as shown in Figure 7 was analyzed. (A) Mass chromatogram of C24:1 on LC-ESI-MS (*m/z* 365.34). The analytical time point is indicated by the arrowhead. (B) Two C24:1 isomers with double bonds either at *ω*-7 or *ω*-9 were discovered. (C) Tandem mass spectra and structures of diagnostic ions at the indicated time point in (A) are shown. Diagnostic fragment ions are indicated by asterisks in the graph.

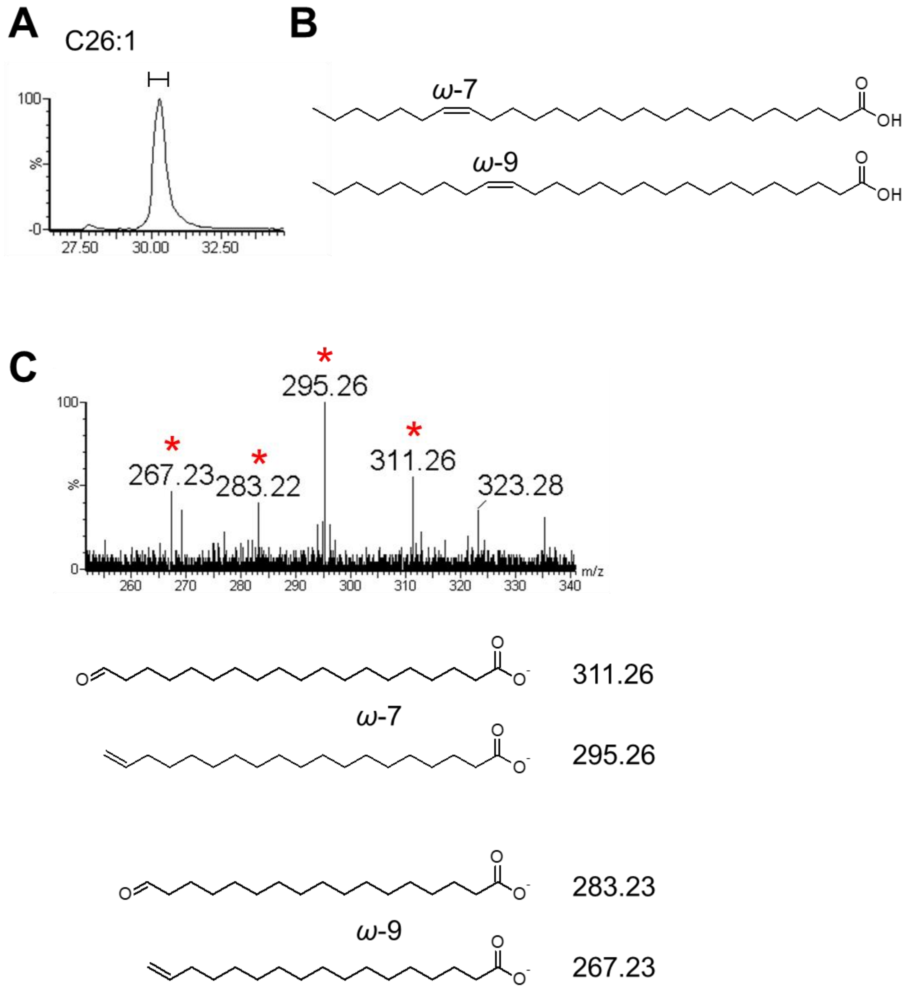


**Figure S11.** Identification of C26:1 FA isomers from human fibroblasts. The same FA sample as shown in Figure 7 was analyzed. (A) Mass chromatogram of C26:1 on LC-ESI-MS (*m/z* 393.37). Because of the low abundance of C26:1 species in the sample, the spectral data were combined throughout the peak, as indicated by the line. (B) Two C26:1 species with double bond either at *ω*-7 or *ω*-9 position were discovered. (C) Tandem mass spectra and structures of the diagnostic ions are shown. Diagnostic fragment ions are indicated by asterisks in the graph.

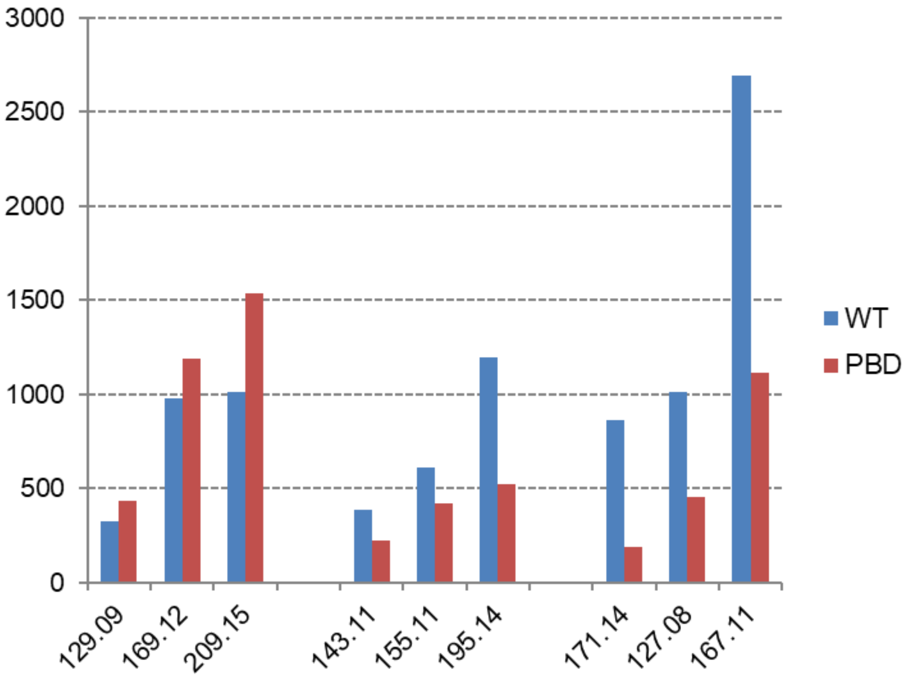


**Figure S12.** Comparison of C20:3 isomers found in human fibroblasts. Intensities of diagnostic fragments from three C20:3 isomers were compared (also see Figure 10). The blue bars represent wild-type fibroblasts and the red bars represent PBD fibroblasts. The numbers on the *x*-and *y*-axes indicate m/z values of diagnostic fragment ions and intensity, respectively.
